# NO modulates human airway smooth muscle function by altering glucose-6-phosphate dehydrogenase effects on sGC function in asthma

**DOI:** 10.64898/2026.05.06.723287

**Authors:** Arnab Ghosh, Mamta P. Sumi, Cynthia Koziol-White, Blair Tupta, Ling Wang, Chaitali Ghosh, William F. Jester, Reynold A. Panettieri, Dennis J. Stuehr

**Author notes:** Both authors contributed equally (Arnab Ghosh, Mamta P. Sumi). Address of Correspondence to: Arnab Ghosh, Department of Inflammation and Immunity/NC22, Lerner Research Institute, The Cleveland Clinic, 9500 Euclid Ave., Cleveland, OH-44196, USA.

## Abstract

Since NO can modulate mesenchymal cell function, we posit that NO can modulate gene expression associated with excitation-contraction coupling. Our study shows that treating asthma-derived HASMCs with a low dose of NO plus sGC stimulator BAY-41, in most cases sensitized smooth muscle sGC towards activation via an elevated sGC heterodimer and in some cases also improved sGCβ1, catalase, Cyb5r3 or Trx1 expression (n=24 non-asthma and n=25 asthma). Interestingly we found that majority of asthma HASMCs showed a marked downregulation of G6PD expression inducing a low GSH/GSSG ratio in asthma, and these findings were replicated in murine lungs of allergic asthma (OVA and CFA/HDM). Studies with HEK/COS-7 cells showed G6PD synergizing with hsp90 in enabling sGC heme-maturation. G6PD overexpression in HASMCs enhanced the sGC heterodimerization while silencing of endogenous G6PD abrogated it. Complementation of these cellular results with whole animal models of G6PD deficiency or overexpression provided verification to our findings. Mouse lung tissue from the humanized variant of G6PD deficiency, V68M (G6PD A-deficiency) showed significant downregulation in the sGC heterodimer, with a concomitant reduction in its NO heme-dependent activity, thereby showing that G6PD deficiency lowers sGC heme. Conversely, G6PD overexpressing mouse lung tissue displayed an elevated sGC heterodimer and also showed a robust G6PD-sGCβ1 interaction, suggesting G6PD to be involved in the heme-maturation of sGCβ1. While G6PD maintains the cell redox by generating NADPH, its new role in regulating sGC maturation links sGC dysfunction in asthma to G6PD deficiency and may potentially uncover new targets for asthma treatment.

## INTRODUCTION

Human airway smooth muscle cells (HASMCs) play a pivotal role in bronchomotor function and in asthma their dysregulation promotes the airway narrowing, obstruction and hypersensitivity that characterizes the disease (1, 2). The β_2_-adrenergic-adenylyl cyclase-cAMP signaling pathway (β_2_AR-sAC-cAMP) that is present in the HASMC modulates bronchomotor tone (3) and is the target of current bronchodilator therapy. HASMC also express soluble guanylyl cyclase (sGC), which by signaling through the nitric oxide (NO)-sGC-cGMP pathway also promotes HASMC relaxation in the small airways (4, 5). The signaling pathways in HASMC that drive bronchodilation can become attenuated in asthma and this typically manifests as defective β_2_-adrenergic receptor activation/recycling (6) or as development of sGC insensitivity toward NO activation (4). Interestingly our studies found that low NO triggers a rapid heme insertion into immature heme-free apo-sGCβ1, resulting in mature sGC heterodimer and that NO can act both ways to make or break the sGC heterodimer (7-9).

Glucose-6-phosphate dehydrogenase (G6PD) is involved in the pentose phosphate pathway, and clinical indicators of primary G6PD deficiency are known to cause hemolytic anemia in response to bacterial/viral infection (10-12), and also contributes to PAH pathogenesis (13-15). G6PD generates NADPH, providing reducing equivalents for antioxidant defense and NADPH oxidase for superoxide generation, and maintains a normal NADPH/NADP ratio that, in turn, regulates glutathione (GSH) biosynthesis. G6PD is classified as a DEG (differentially expressed gene) and studies found that G6PD deficiency is associated with an increased risk of asthma (16-20), warranting further assessment in asthma patients. More evidence of G6PD involvement is supported by recent bioinformatics studies that identified G6PD as one of the 5 hub genes with clinical diagnostic value in asthma (21). In a lung environment, G6PD deficiency may cause the oxidative setting to be high, leading to elevated NOS in the airway epithelia (iNOS) or in the lung endothelium (eNOS), thereby becoming heme-free and diminishing the capacity of NOS enzymes to generate NO. This incapacitates smooth muscle sGC as it becomes heme-free or low in expression and in the absence of its natural activator NO, bronchodilation is hindered. A similar circumstance was revealed by our recent study on PAH (22), where insufficient eNOS activation and low sGC expression caused poor sGC heterodimerization (22), resulting in insignificant dilation to the arteries. Treating PASMCs derived from PAH (hereditary or idiopathic) with a low dose NO produced a significant elevation in the heterodimer leading to increased sGC activation. Low NO thus leapfrogged the hindrances to activate the sGC, and these findings may have similar implications for curing the dysfunctional sGC in asthma. Our earlier studies showed that a majority HASMCs from severe asthma had a unresponsive, dysfunctional sGC characterized by a poor heterodimer and low expression of certain support/redox proteins like Cyb5R3, Trx1 and catalase (23-25). We thus investigated asthma derived HASMCs response to low NO, and tested both asthma HASMCs and murine models for G6PD expression deficiency to assess an association with occurrence of heme-free sGC in asthma. The results from this study suggest that low NO can restore activation of a dysfunctional sGC in asthma and indicates a novel role of G6PD in enabling heme-maturation of sGCβ1.

## RESULTS

### Low NO induces sGC heterodimerization in PCLS and asthma HASMCs

Our earlier studies found that sGC agonists and NO can dilate small airways in PCLS (5). In order to determine the impact of low NO on sGC activation and its associated molecular signatures in human PCLS, NO doses from DETANONOate (0-10 μM) were used to treat PCLS for 4, 18 and 24 h. As depicted in Fig. 1, the basal levels of sGC activation by low NO on the PCLS peaked at 10 μM for both time points (4 & 24 h), and was also reflected in the BAY-41 response relative to BAY-60 (Fig. 1A). Immunoprecipitation (IP) assays done on tissue supernatants to assess molecular signatures of sGC activation showed a gradual increase in sGCα1β1 heterodimerization with a concomitant fall in sGCβ1-hsp90 interactions (Fig. 1B) (23), suggesting low NO activates sGC in the small airways by increasing its heterodimer (7, 22). These results and our prior studies on NO restoring vascular sGC in PAH PASMCs (22), prompted us to study the impact of low NO on HASMCs derived from severe asthma (Table 1 and Fig. S1). HASMCs were first cultured to assess expression of sGCβ1 or certain redox enzymes which may impact the heme-dependent sGC activation (23). As depicted by western blots in Fig. S2, we found greater expression of sGCβ1, and low mean expression of sGCα1, along with certain redox proteins like catalase or Trx1 in asthma relative to non-asthma donors (Fig. S2A). This was similar to an earlier study (n=16, asthma HASMC) (23) which displayed a similar pattern of expression and majority of cells showed an inherently defective sGC phenotype that was heme-free and BAY-60 responsive. As depicted in Fig. 2A, a low dose of DETANONOate for 16 h plus 10 min of BAY-41 treatment, induced sGC heterodimerization and subsequent activation in 8/8 non-asthma or 6/8 asthma HASMCs (Set I, Fig. 2A, B). In the other 6 non-asthma or asthma HASMCs from set I, only 2 asthma cells were unresponsive to low NO (Fig. S2B). This implied that low NO doses could restore sGC function in over 70% of asthma-derived cell lines (10/14 in Set I), and it activated all the non-asthma-derived HASMCs. The results were more pronounced in the next set of 11 asthma HASMCs, as over 90% (10/11 HASMCs) were activated by low NO + BAY-41 (Set II, Figs. 2C, D). All the non-asthma cell lines were activated in this set (Set II, Figs. 2C, D) and the activation response coincided with an increase in the sGC heterodimer in all cells (Sets I & II, Figs. 2A-D). The asthma HASMCs which failed to respond to the low NO + BAY-41 remained BAY-60 responsive (Sets I & II, Figs. 2C & D). Together, our data suggests that low NO activates sGC in human PCLS and asthma-derived HASMCs.

**Figure 1.**
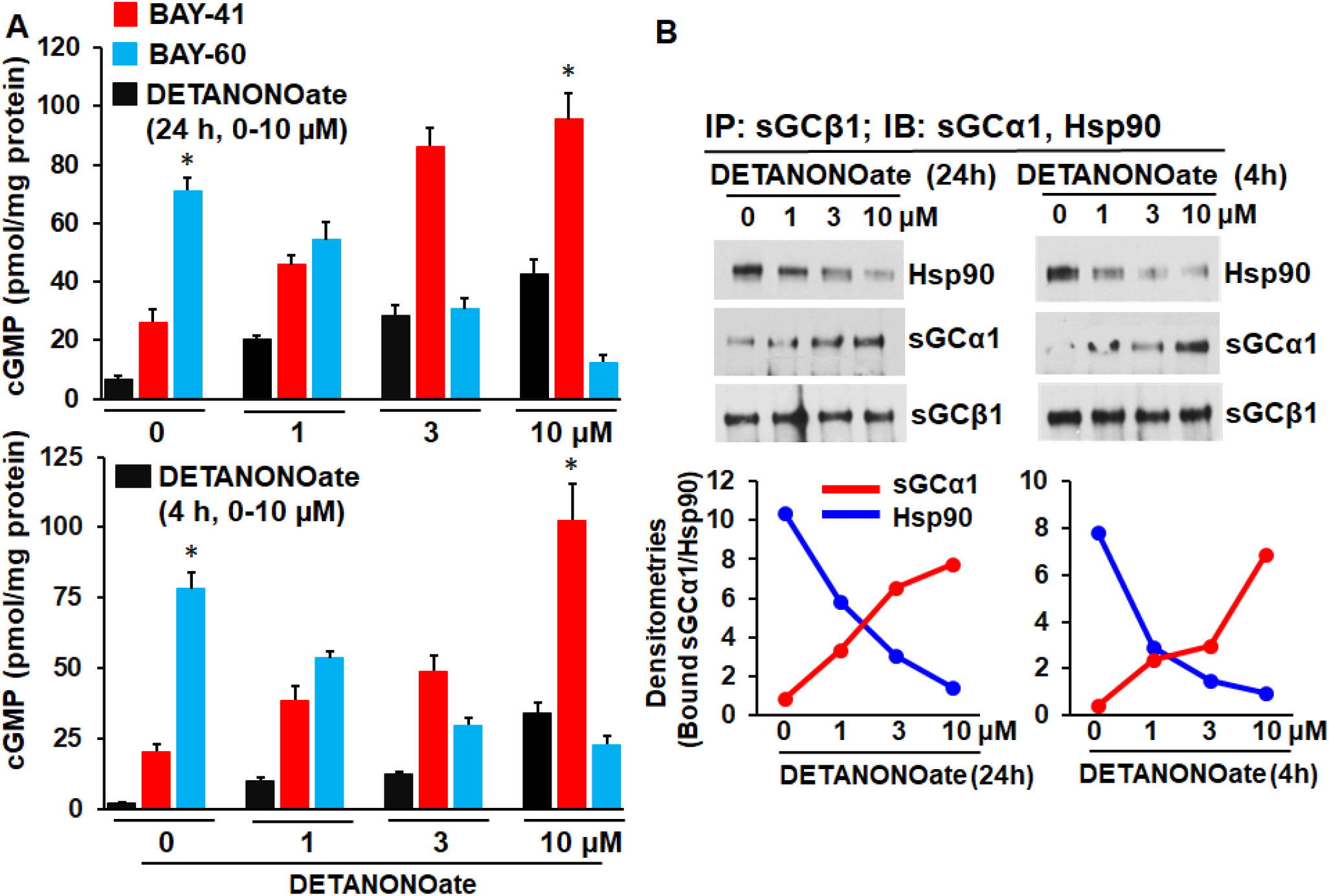
Impact of low NO from DETANONOate on sGC activation in PCLS. PCLS were treated with low NO doses (0-10 µM, 4 or 24 h), and the impact of NO on sGC activation and its associated molecular signatures analyzed. PCLS supernatants were used to assess sGC activation and IP assays to test sGCα1/Hsp90-β1 interactions. (A) cGMP estimates to assess sGC activation following stimulation with BAY-41/60 on PCLS pretreated ± low NO. (B) Corresponding IP assays to assess sGCα1/Hsp90 bound to sGCβ1 and densitometries of the immunoblot bands. Values are mean (n=3 repeats) ± SD. *p < 0.05, by student’s unpaired *t*-test. Bars marked by * are significant.

**Figure 2.**
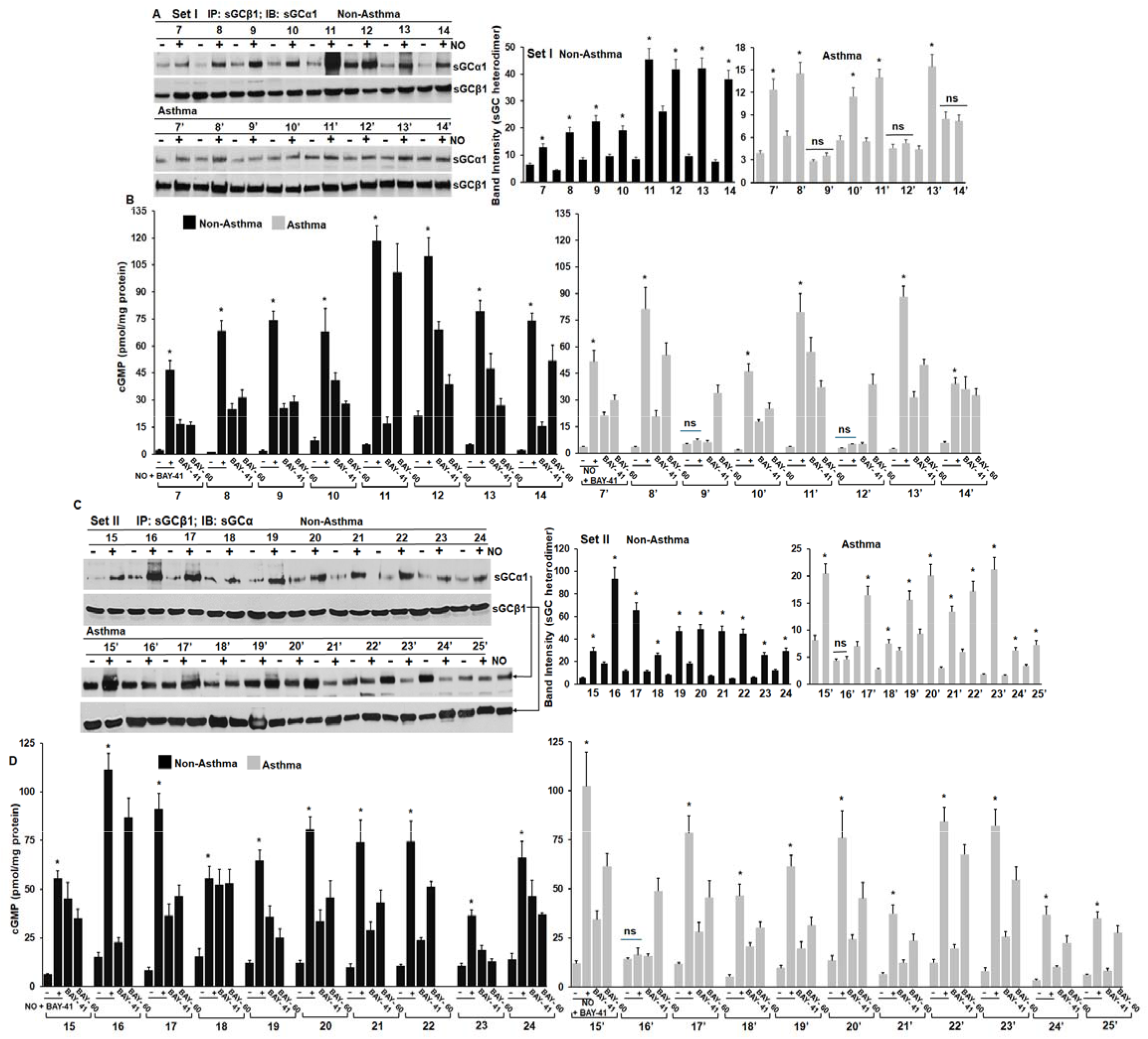
Impact of low NO on sGC heterodimerization and activation on asthma derived HASMCs. HASMCs from non-asthma or asthma patients were cultured to confluency and then treated with a low dose of DETANONOate (5 μM, 16 h), followed by BAY-41 treatment before cell harvest while parallel cultures were untreated or treated with BAY-41/60. (A & C) IP assays following low NO treatment on normal or asthma HASMCs. An increase in sGC heterodimer with NO corresponds to subsequent sGC activation. (B & D) sGC activation assessed by cGMP generation. Calculated densitometries of sGCα1 bound to sGCβ1 are mean (n=3 repeats) ± SD. *p < 0.05, by student’s unpaired *t*-test. Bars marked by * are significant relative to the non-NO treated HASMC. ns is statistically non-significant.

### Non-responder asthma HASMCs seem inclined towards a desensitized sGC phenotype

In our earlier study (23) we found that sGC was inherently defective in most of the asthma HASMCs (12 out of 17 or over 70%), where these cells were primarily BAY-60 responsive (>50% of the asthma HASMCs in set 1 were BAY-60 responsive) (Fig. 2B). In order to restore sGC activity in these asthma HASMCs, we treated these cells with a low dose of NO + BAY-41. While some of asthma HASMCs did not respond to low NO (7/10 from the prior set{Fig. S3} and 2/8 or 1/11 asthma HASMCs in Set I or II respectively {Fig. 2}) nor did their sGC heterodimer increase during the NO treatment, and these can be categorized into responders and non-responders. These suggest that the non-responder cells may be genetically predisposed towards a desensitized sGC phenotype.

### Low NO induces sGC or redox protein expression in some asthma HASMCs that may not necessarily correspond to sGC activation

In the current study some HASMCs from both non-asthma/asthma (Figs. 3 and S4A) showed induction of expression for sGCβ1, catalase, Cyb5r3, Trx1 or G6PD but not sGCα1, suggesting that NO can modulate certain genes at transcriptional levels in addition to having an impact on sGC heterodimerization (Figs. 2, 3, 4 and S4A). While expression of sGCβ1 was equivocal in asthma HASMC relative to non-asthma in the current sets, as it was variable for asthma HASMC in sets 1 and II, sGCα1 expression was variable in Set I but more uniform in Set II (Figs. 3, S2A, S4A & table S3). Variations in expression were also found for catalase, Cyb5r3, Trx1 or G6PD in both Sets (Figs. 3, S2A, S4A & table S3). On selected HASMC lines we used specific primers to evaluate mRNA expression of *sGC*β*1, Cyb5r3* or *Trx1* −/+ NO, and found that the mRNA expression data supports the protein data showing induction of these proteins (Fig. S4B & table S3). Our analysis of the data depicted in Figs. 3, S3, and Table S3 suggests that for proteins that were low in expression, like G6PD or sGCβ1, 6/9 asthma HASMC lines showed increased expression for G6PD, and 2/8 for sGCβ1, following low NO that also led to increased sGC activation. Likewise, it was 2/7, 8/8, or 2/8 asthma HASMCs showed increased catalase, Trx1 and Cyb5r3 expression, respectively. Here, low NO differentially induced protein expression in some but not all asthma HASMCs. Together, these suggest that modulation by low NO differentially impacts HASMCs, and asthma HASMCs 2’, 3’, 9’, 12’ and 16’ (sets I and II) can be categorized as non-responders because despite protein induction by low NO, their sGC could not be activated (Table S3 and Fig. S4).

**Figure 3.**
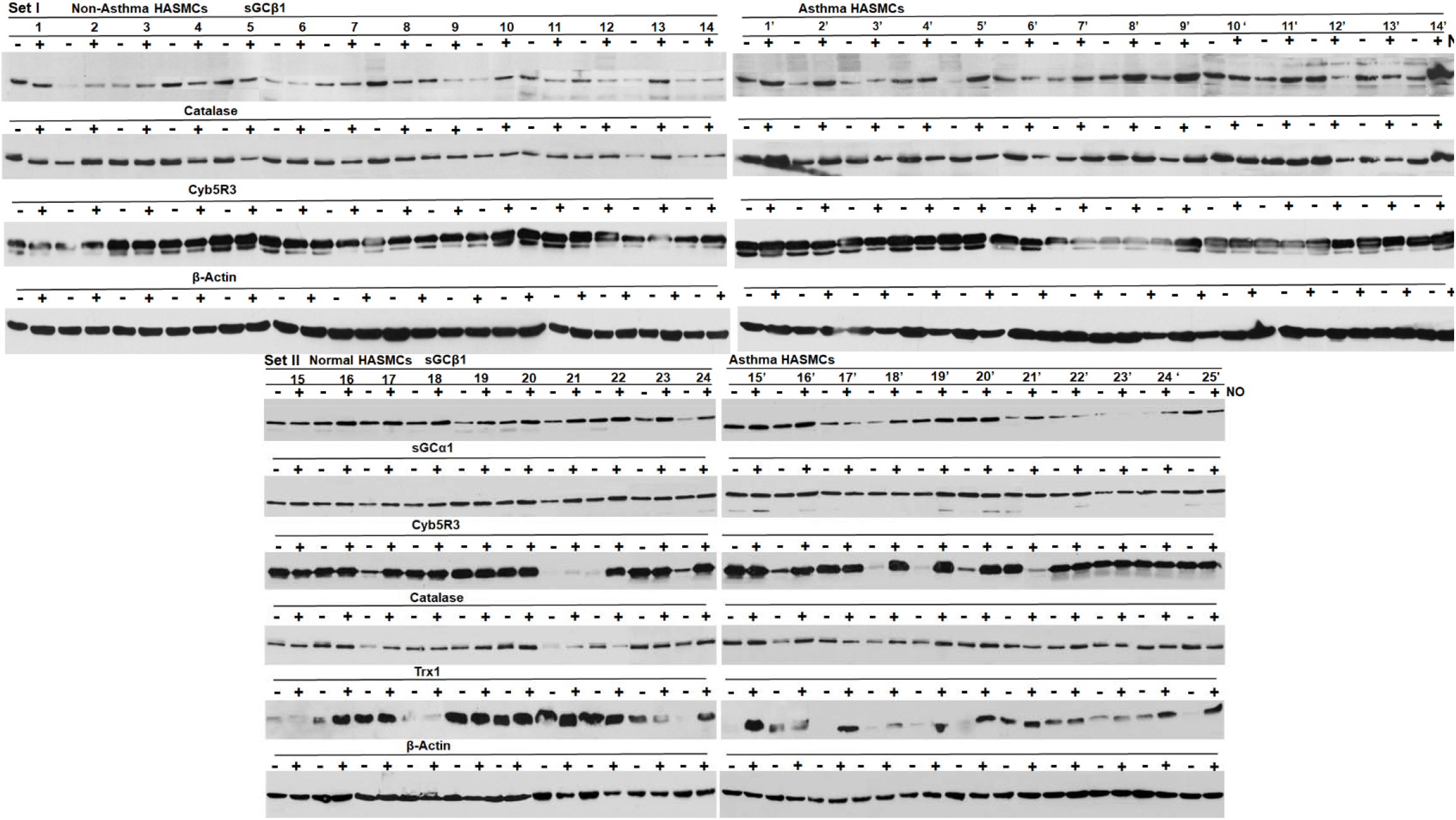
sGC and redox protein expression and its modulation by low NO. Supernatants prepared from non-asthma or asthma HASMCs that were treated with DETANONOate (16 h) and western blotted with specific antibodies. (A & B) Expression levels as indicated from Set I or Set II HASMCs.

**Figure 4.**
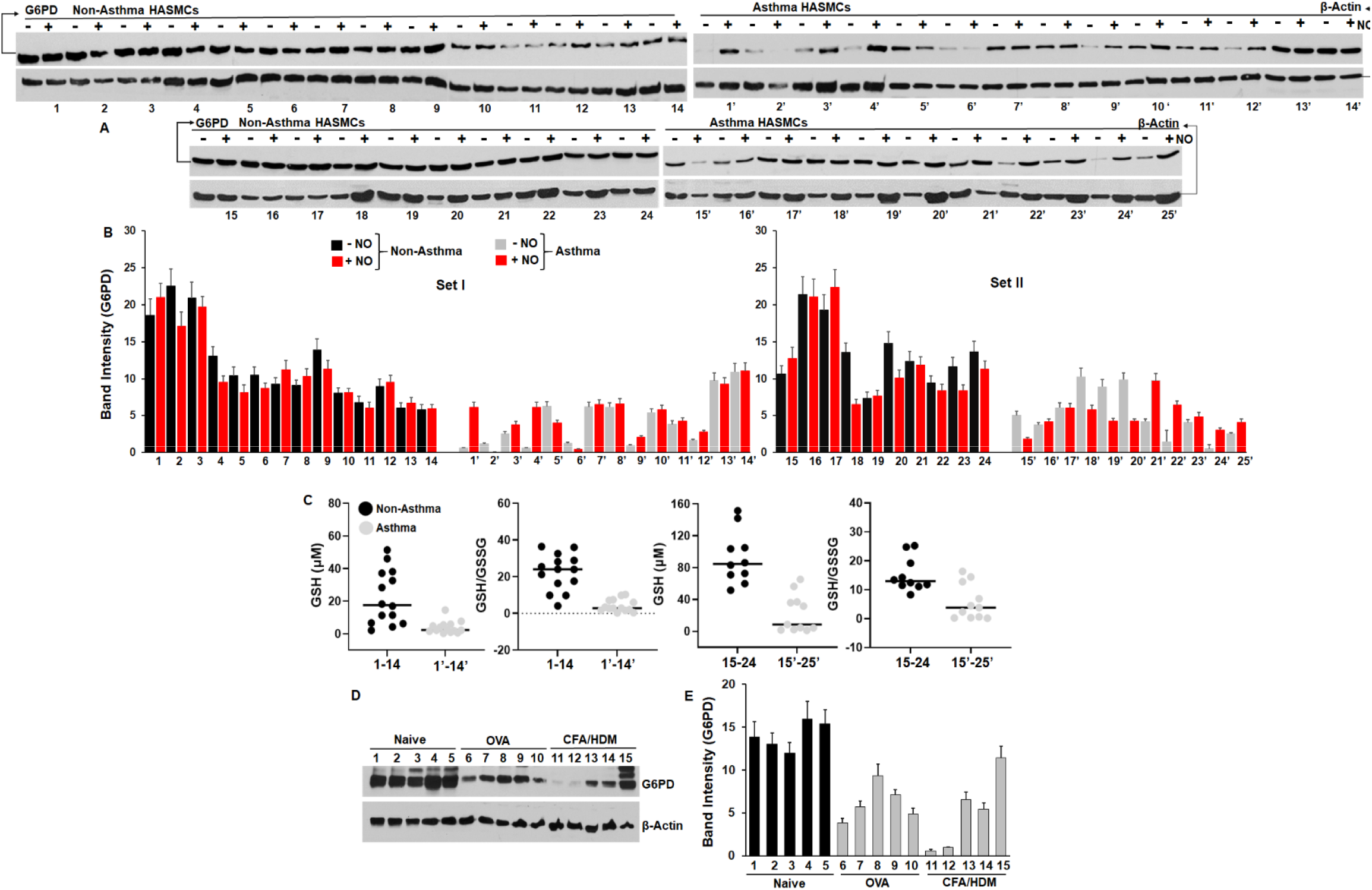
Expression of G6PD and its corresponding GSH/GSSG ratios in HASMCs or in lungs of murine asthma. HASMC from non-asthma or severe asthma patients were cultured to confluency and treated with a low dose of DETANONOate (5 μM, 16 h) before cell harvest. The cell supernatants or those from mouse lungs of control naïve or asthma models; OVA and CFA/HDM (n=5 lungs for each) were western blotted with G6PD and β-actin antibodies. Additionally, GSH/GSSG measures were made on HASMC supernatants. (A & D) Western blots for protein expression as indicated. (C) GSH and GSH/GSSG measures as indicated. (B & E) Corresponding mean (n=3 repeats) densitometries calculated from band intensities as depicted in panels A or D normalized to β-actin used as a loading control.

### G6PD expression is downregulated in asthma

As G6PD is known to be a risk factor in asthma and deficiencies in G6PD are associated with development of PAH (13-15), we sought to determine whether low expression of G6PD conferred a highly oxidative milieu that would eventually contribute to sGC becoming heme-free in asthma. As depicted in Fig. 4, we found markedly lower expression of G6PD in most of the asthma HASMCs relative to the non-asthma, with majority asthma HASMCs being low in G6PD expression (Fig. 4A, B). This was also the case for majority of asthma HASMC relative to non-asthma used in our previous study (23) (Fig. S3). Moreover, fewer HASMC lines showed significant induction in G6PD protein levels following stimulation with low NO (none for the non-asthma and 5/14 in Set I or 4/11 in Set II for asthma HASMC, or 9/49 total cells, Fig. 4B). Assaying for GSH expression or GSH/GSSG ratio in asthma HASMC showed a low GSH/GSSH ratio in the asthma HASCM relative to the non-asthma HASMC (Fig. 4C) and this low G6PD expression corroborated with its low activity (GSH/GSSH ratio, Fig. S5). The low expression of G6PD in these human asthma cells was similar to expression studies from murine models of allergic asthma, as lung tissues from OVA and CFA/HDM models also showed downregulation in their G6PD levels (Fig. 4B-E). Together, these results demonstrate that cellular G6PD levels are downregulated in asthma HASMC, resulting in a low GSH/GSSG ratio in these cells.

### G6PD synergizes with hsp90 to promote sGC heterodimerization that is elevated by G6PD overexpression or diminished by its silencing

To determine the impact of G6PD on sGC heterodimerization and activation, we overexpressed G6PD by transient transfection in COS-7 cells along with sGCα1β1 constructs. Following transfection (42 h), cells were treated with low dose of NOC-18 for 16 h before BAY-41 treatment (10 min) and cell harvest. Transient G6PD expression induced 3 fold more expression of G6PD over the endogenous G6PD (Fig. 5A). As depicted by IPs in Fig. 5B, we found that G6PD interacted with sGCβ1 and hsp90 prior to NO-induced activation, but the interaction dissociated with NO activation when the subunit heterodimerization peaked. There was a stronger heterodimer formed (over two-fold) with G6PD overexpression relative to non-transfected cells, with enhanced sGC activation in the G6PD overexpressing cells, as determined by cGMP production (Figs. 5B, C). We further assessed the G6PD-sGCβ1 interaction in the unactivated state (-NO) by immunostaining using non-asthma/asthma HASMCs, and found that these co-localized (Fig. S6). These indicated that G6PD, like hsp90, may play a role in sGCβ1 heme-maturation. To test this idea, we expressed sGCα1β1 in SA pretreated (48 h) COS-7 cells and determined sGCβ1 interaction with G6PD under heme-depleted (+SA) or heme-repleted (+SA +hemin for 3h) conditions relative to non-SA controls (-SA). Using IP assays (Fig. 5D), we found G6PD-hsp90-sGCβ1 association under normal conditions (-SA) (Fig. 5E), indicating the presence of pre-existing heme-free sGCβ1 (8). More importantly endogenous G6PD synergized with hsp90 and associated with the heme-free sGCβ1 under +SA conditions, and these interactions dissociated post heme-insertion (+SA+hemin, Fig. 5E) (8, 26). These data show increased sGCα1β1 heterodimer induced by the presence of G6PD (Fig. 5B), and that the enhanced binding of G6PD to heme-free sGCβ1 (Fig. 5D) was unperturbed even in the face of low G6PD levels that may develop under heme-free conditions (+SA, Fig. S7). To verify these results, we transfected G6PD into HEK cells along with another protein Mb, whose heme-insertion/maturation is also hsp90-dependent (27). As depicted in Fig. 5F-H, we found that G6PD overexpression enhances sGC heterodimer formation, while co-expression of G6PD with Mb appeared to reduce the sGC heterodimer during NO activation, as the equilibrium may shift the towards heme maturation of Mb, which like sGC, is also hsp90-dependent (27). Similar results were obtained by Mb expression in HEK cells, as it reduced the endogenous sGC heterodimer (Fig. 5G, H). These results show that G6PD synergizes with hsp90 to enable heme-insertion into sGCβ1, causing subsequent heterodimer formation. To test for G6PD promoted sGC heterodimerization in primary HASMCs we overexpressed G6PD in 4 non-asthma and 7 asthma HASMCs. This resulted in increased basal levels of sGC heterodimer (-NO, unactivated) and induced significant heterodimer formation during activation (+NO) as shown by increased cGMP levels (Figs. 6A-C).

**Figure 5.**
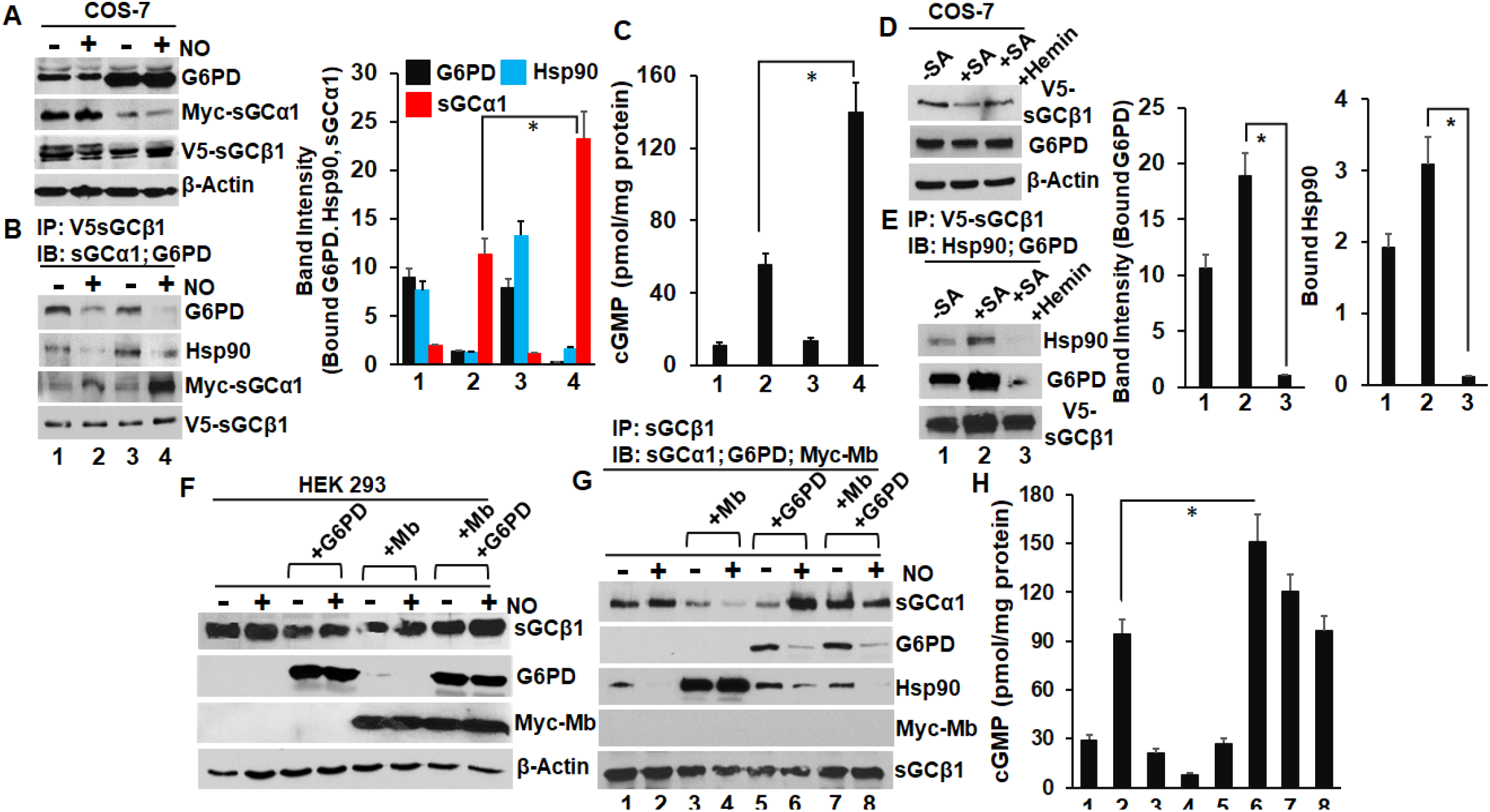
Hsp90 and G6PD synergizes with heme-free sGCβ1 during its heme-maturation and impact of G6PD on sGC heterodimerization. COS-7 cells transfected with sGCα1β1 and/or G6PD constructs (36 h), then treated with a low dose of NO donor from NOC-18 for 16 h before BAY-41 (10 min) treatment and cell harvest. Parallel experiments were performed by making these cells heme-free by pre-treating them with SA (for 48 h), followed by transfection of sGCα1 & β1 for additional 42 h before hemin addition (3 h) and cell harvest. In separate experiments HEK 293T cells were either untransfected or transfected with Mb and/or G6PD (36 h), followed by low NO (from NOC-18 for 16 h) + BAY-41 treatment and cells harvested. The generated cell supernatants were analyzed for protein expression by western blots, protein-protein interactions by IPs or sGC activation by cGMP estimation. Panels (A, D & F) Representative western blots depicting protein expression as indicated and its mean densitometries from n=3 repeats, ± SD. Panels (B, E & G) IP assays depicting the impact of G6PD or Mb overexpression on sGCα1β1 heterodimerization or the impact of expressing heme-free sGC on sGCβ1-G6PD/hsp90 interactions. Densitometries calculated from band intensities of sGCα1-β1 or sGCβ1-G6PD/hsp90 interactions are mean (n=3 repeats) ± SD. For panels A-E, *p < 0.05, by student’s unpaired *t*-test. Panels (C & H) cGMP estimation by ELISA. Depicted values are mean of n=3 repeats ± SD. *p < 0.05, by one-way ANOVA, followed by post-hoc analysis based on a *t*-test with Bonferroni adjustment where *p < 0.01667 and all conditions were significantly different in each case.

**Figure 6.**
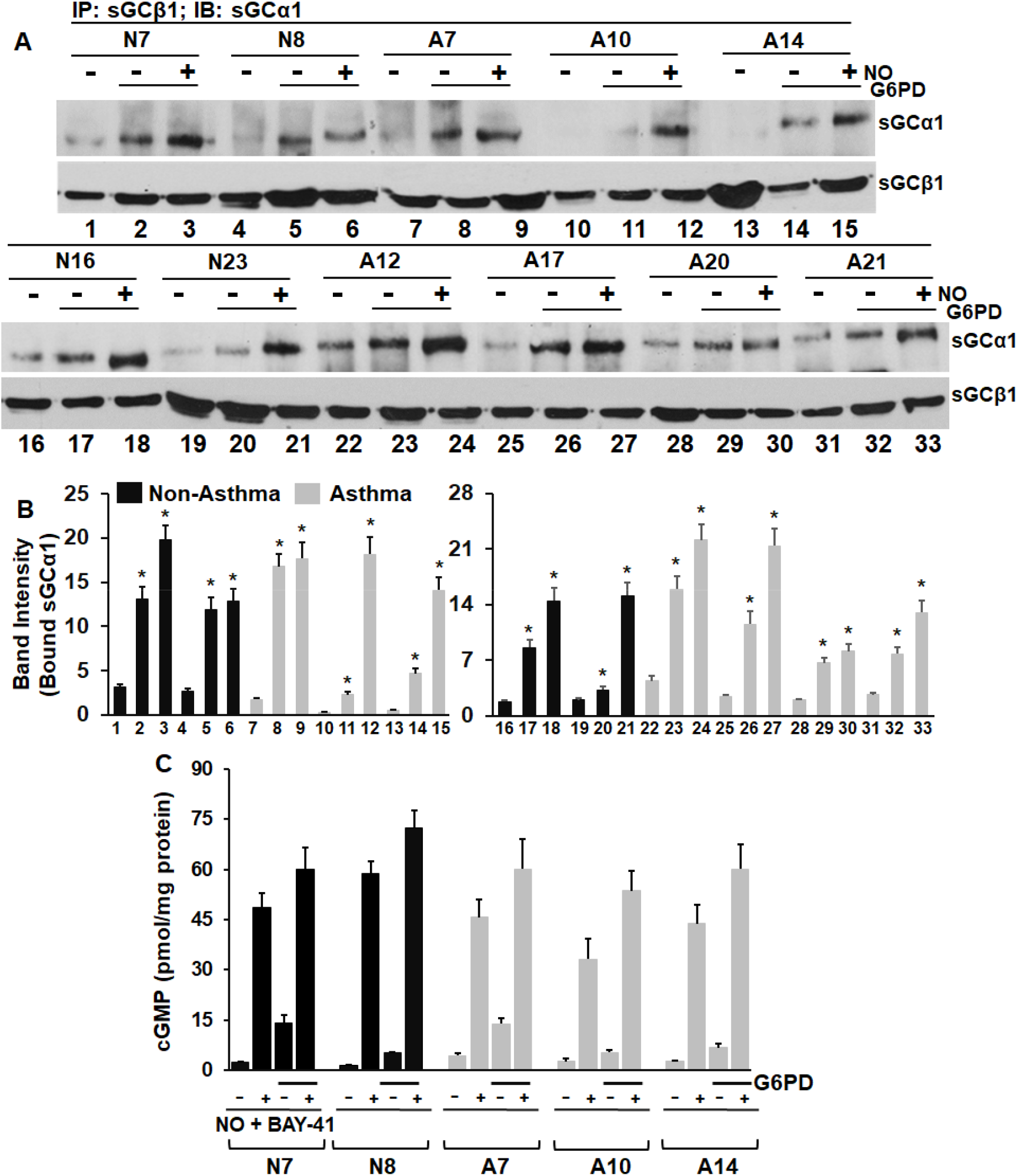
Impact of G6PD overexpression in HASMCs on sGC heterodimerization and activation. To determine whether G6PD in promotes sGC heterodimerization in primary cells (four non-asthma and seven asthma HASMC) we overexpressed G6PD by transient transfection (42 h) followed by a dose of low NO from DETANONOate for 16 h + BAY-41 before cell harvest. The supernatants were analyzed for protein-protein interactions by IPs or sGC activation by cGMP estimation. (A) IP assays depicting the impact of G6PD overexpression on sGCα1β1 heterodimerization. Densitometries calculated from band intensities of sGCα1 bound to sGCβ1 are mean (n=3 repeats) ± SD. (B) cGMP estimation by ELISA. Depicted values are mean of n=3 repeats ± SD. *p < 0.05, by student’s unpaired *t*-test.

We then reduced the endogenous G6PD levels in HASMCs with G6PD-targeted siRNA. Transfecting G6PD siRNA (48 h) significantly attenuated endogenous expression of G6PD (~2-5 fold downregulation, Fig. 7A) relative to control scrambled siRNA and sGC heterodimerization was also significantly reduced by ~4-6 fold during NO stimulation (Fig. 7B), with subsequent reduction in the cGMP levels (Fig. 7C). Together, these results suggest that G6PD plays critical role in enhancing sGC heterodimer formation in HASMCs, and this role of G6PD appears to be distinct from its role in NADPH generation or maintaining the GSH/GSSG balance in cells.

**Figure 7.**
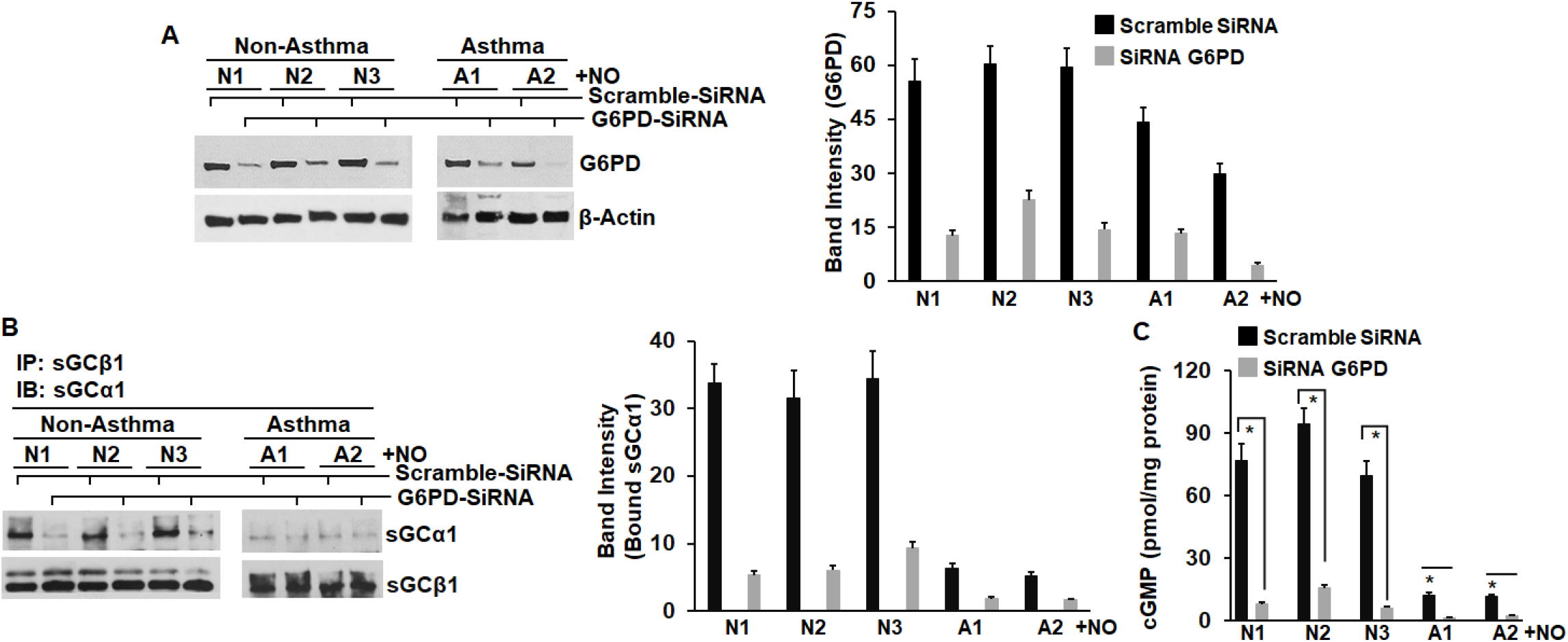
Silencing of endogenous G6PD in HASMCs abrogates sGC heterodimerization and its subsequent activation by NO. To study the impact of endogenous G6PD on sGC heterodimerization and activation, we transfected the siRNA designed against human G6PD for 48 h, followed by low NO treatment for 16 h in select HASMCs (three non-asthma and two asthma) and compared the results relative to control scrambled siRNA transfections. The generated supernatants were analyzed for protein expression by immunoblot, protein-protein interactions by IPs or sGC activation by cGMP estimation (A) Representative western blots depicting G6PD expression as indicated and its mean densitometries from n=3 repeats, ± SD, normalized to β-actin. (B) IP assays depicting the impact of endogenous G6PD silencing on sGCα1β1 heterodimerization. (C) cGMP estimation by ELISA. Depicted values are mean of n=3 repeats ± SD. *p < 0.05, by student’s unpaired *t*-test.

### Modulation of G6PD activity impacts sGC function in murine models

To determine the impact of G6PD deficiency on lung sGC function we used tissues from a novel mouse strain carrying the “humanized” African G6PD variant, V68M (G6PD A-). This represents the most common G6PD variant displaying ~5% G6PD activity relative to the WT (28). As depicted in the Fig. 8A, although there was equivalent expression of sGCβ1 in the lung tissue supernatants of G6PD V68M and WT, the V68M mice displayed a significantly reduced sGC heterodimer, that was > 2.5 fold reduced relative to the WT mice (Fig. 8B). This resulted in a lower heme-dependent sGC activation by NO or BAY-41 for the V68M variant relative to WT (Fig. 8C). As the G6PD activity in V68M mice is ~90% lower relative to the WT mice, these data indicate that lowered G6PD activity of this variant severely impacts the sGC function. We then compared these findings using lung tissue supernatants from G6PD overexpressing transgenic mice that has been shown to improve median lifespan in female mice via higher NADPH levels and provides better protection from the deleterious effects of ROS (29). These mice exhibited moderately enhanced lung G6PD expression (Fig. 8D), while expression of sGCβ1, its heme-support chaperone hsp90, or redox enzymes like Trx1 were comparable to levels in WT mice. IPs of lung supernatants showed that in G6PD overexpressing mice, the sGC heterodimer was ~3 fold elevated, while G6PD-bound sGCβ1 was elevated >33% relative to WT (Fig. 8E). There was also a linear correlation between sGCβ1-G6PD interactions and sGC heterodimer, suggesting that G6PD may participate in the heme-maturation of sGCβ1 (Fig. 8E). This was further confirmed by sGC activation assays that showed a significant increase in the heme-dependent sGC activity by NO or BAY-41 in lung supernatants from G6PD mice relative to WT (Fig. 8F). We also found a linear correlation between G6PD expression and HASMCs response to low NO (Fig. S3). Together these findings suggest that G6PD may be a novel regulator of sGC heme.

**Figure 8.**
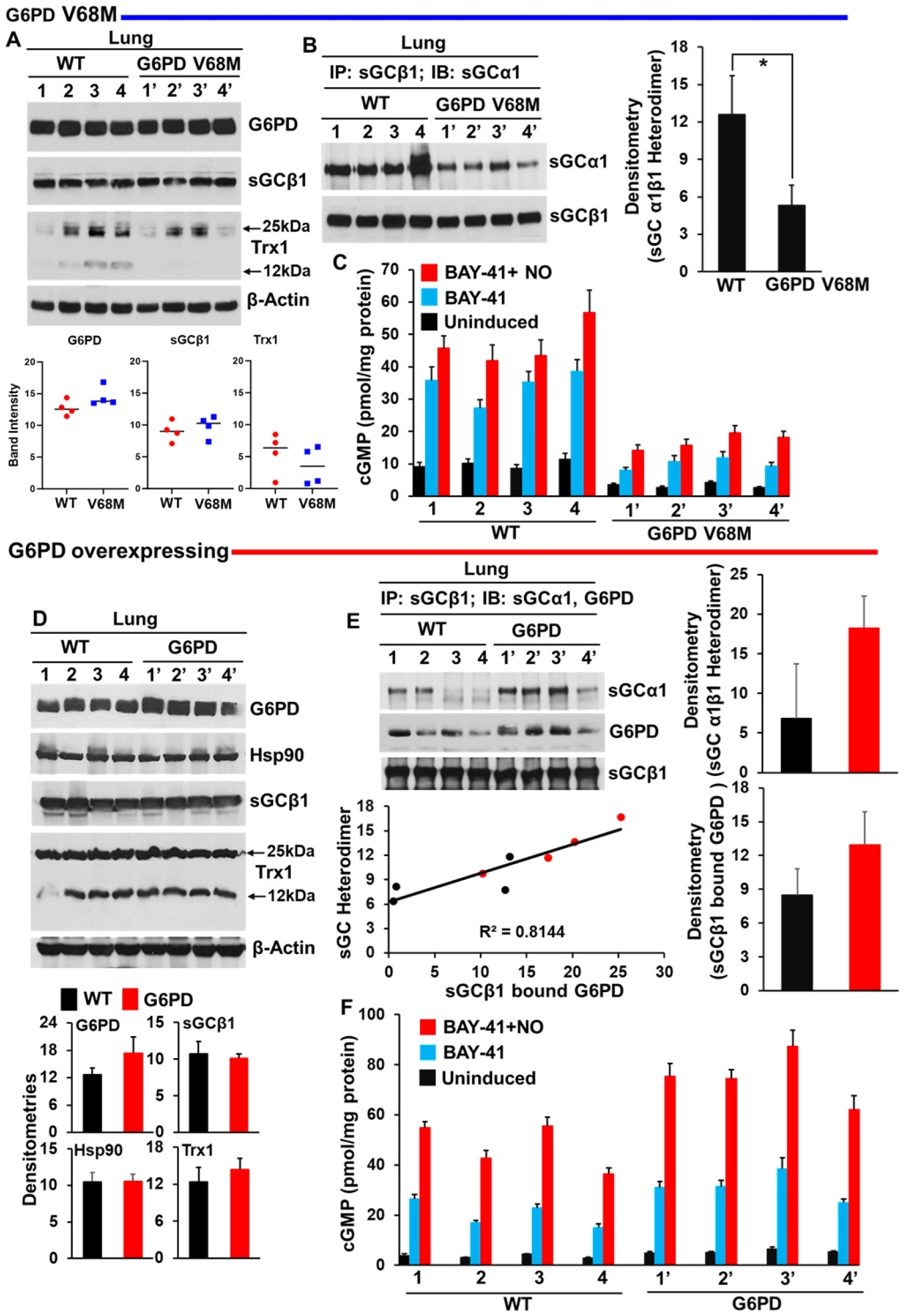
Impact of G6PD (V68M) variant or G6PD overexpressing transgenic mice on lung sGC heterodimer and activity. Lung supernatants from G6PD V68M male hemizygous mice (n=4) or from G6PD overexpressing transgenic female mice (n=4) were subjected to expression analysis by western blots, IPs and activation assays by sGC stimulator BAY-41. (A & D) Protein expression of indicated proteins with corresponding mean (n=4) densitometries. (B & E) IP assays depicting the impact of G6PD V68M or G6PD overexpression on sGCα1β1 heterodimerization relative to WT. Densitometries calculated from sGCα1 bound to sGCβ1 are mean values, −/+ SD from n=4 mice lungs. For panel E, Correlations between individual desitometries calculated from sGCα1 bound to sGCβ1 and sGCβ1 bound to G6PD are also depicted along with the correlation coefficient. *p < 0.05, by an independent student’s *t*-test. (C & F) cGMP estimation by ELISA as a read out for sGC activation by NO and or BAY-41. Depicted values are mean of n=3 repeats ± SD.

## DISCUSSION

Our current study showed that we are able to restore NO-desensitized sGC in asthma HASMCs, and in most cases, that there was a heterogeneity with respect to restoration of this response in the asthma HASMC lines tested. This restoration was achieved with low NO treatment through elevation of the levels of sGC heterodimer, and in certain cases, through the expression of sGCβ1, catalase, Cyb5r3 or Trx1 (Figs. 2, 3, S2 and S4). The modulation by low NO differentially impacted HASMCs, and certain asthma HASMCs can be categorized as non-responders because despite protein induction by low NO, their sGC could not be activated and these non-responders seemed inclined towards a desensitized sGC phenotype. (Table S3 and Figs. 2, S3, S4). Our findings also show that a majority of asthma HASMCs, as well as murine models of allergic asthma (OVA and CFA/HDM), were associated with a significant downregulation of G6PD expression (Fig. 4A-D, S3 & S5), which were verified by low GSH concentration and a low GSH/GSSG ratio in asthma HASMC relative to non-asthma HASMC (Figs. 4C, S5). These data suggest that high oxidant stress causing oxidation of the ferrous sGC heme to its ferric form maybe associated with asthma HASMCs and eventually contribute to sGC becoming heme-free (4, 23). Accordingly, the G6PD downregulation may thereby contribute to the buildup of heme-free sGC in HASMC, or in murine asthma models (4, 23, 30). Our studies with HEK or COS-7 cells showed that G6PD is likely involved in the maturation of sGC, as G6PD bound to heme-free sGCβ1 in SA treated cells, synergized with hsp90, and positively impacted sGC heterodimerization (Fig. 5 and S7). G6PD overexpression in non-asthma or asthma HASMC enhanced sGC heterodimerization, while silencing of endogenous G6PD abrogated sGC heterodimerization and its subsequent activation by nitric oxide (NO) (Figs.6 and 7). Complementation of these cellular results with whole animal models of G6PD deficiency or overexpression provided verification of our HASMC data. Mouse lung tissue from the humanized variant of G6PD showed significant downregulation in the sGC heterodimer, and showed a concomitant reduction in its NO heme-dependent activity relative to tissues from WT mice, thereby showing that G6PD deficiency lowers sGC heme (Fig. 8A-C). Conversely, G6PD overexpressing mouse lung tissue displayed an elevated sGC heterodimer, which showed a robust G6PD-sGCβ1 interaction, suggesting G6PD to be involved in the heme-maturation of sGCβ1 (Fig. 8D-F). While cellular G6PD maintains the redox milieu in cells by generating NADPH and keeping the GSH/GSSG ratio in the balance (16, 17, 29, 31, 32), its role in regulating sGC heme maturation is a new and novel finding. Our data suggest that this new role of G6PD can add it to the repertoire of metabolic enzymes exhibiting non-metabolic functions (33). Our finding sheds light on a role of G6PD as a regulator of lung sGC maturation, connects sGC dysfunction to its de-activation in human asthma to deficient G6PD expression, which thereby compromises the NO-sGC-cGMP signaling axis in asthma.

G6PD provides reducing equivalents to NADPH oxidase, but also reduces ROS or causes ROS scavenging that is considered beneficial to cell function (34). Hence, G6PD activity modulation has been shown to produce both harmful as well as beneficial effects in different inflammatory conditions either through its inhibition or overexpression (29, 35, 36). As NADPH is required for the reduction of ROS, G6PD deficiency usually results in increased inflammation and cell damage (35, 37-39). G6PD deficiency in cells causes increase in inflammatory molecules or can increase cell adhesion molecule expression (40, 41). G6PD knockdown in HepG2 cells induced IL-8, and tissues that have low G6PD expression have been shown to be more prone to oxidant□induced injury (42). Therefore, overexpression of G6PD has been shown to be protective in different organs (32, 36), with G6PD overexpressing mice displaying an extended median lifespan (29) and a more stable sGC heterodimer (Fig. 8). However, under certain inflammatory conditions such as atherosclerosis, heart failure and obesity, increased availability of NADPH from G6PD has been shown to promote NOX□derived ROS, that over titrates its neutralization leading to increased inflammation (43-45). Likewise G6PD activity has been shown to be elevated in animal models of acute lung injury (46) and in this scenario, pharmacological inhibition of G6PD may be beneficial due to the limited supply of NADPH. In G6PD deficiency, as in asthma, the decreased supply of NADPH limits GSH regeneration, lowering the GSH/GSSG ratio and, in turn, hinders the disposal of oxidants (47). The NADPH produced in the PPP is also a substrate for NOSs to generate NO from L-arginine and for NOX to release the superoxide anion (16). A G6PD deficiency thereby causes reduced neutralization of superoxide or other free radicals and also causes NO depletion (48) that may impact bronchodilator tone. Therefore, G6PD deficiency causing a high oxidative environment with low NADPH or GSH levels can make both NOSs and sGC heme-free, hampering vasodilation in PAH or bronchodilation in asthma. G6PD was downregulated in asthmatic children (19, 20) with low G6PD activity in patients with atopic asthma (49), and a study found a credible association between asthma and G6PD deficiency (17). In another study GEO databases (21) were used to retrieve asthma-related datasets, where bioinformatic studies identified 1698 DEGs linked to asthma, and G6PD was identified as one of the 5 hub genes with clinical diagnostic value in asthma.

While studies on G6PD expression on airway smooth muscle are scanty our studies explicitly show that significantly low levels of G6PD expression and activity occurs in severe asthma (Figs. 4, S3) that can solely drive sGC towards a desensitized heme-free phenotype. Increase in G6PD levels are also reported in bronchial epithelial cells in asthma (21) and this may support high induction of iNOS or NO production via high NADPH levels, that eventually inactivates the smooth muscle sGC (4). So both high G6PD levels supporting high NO in the airway epithelia or its deficiency in the smooth muscle may downregulate sGC function (7). The clinical significance of our study lies in selectively boosting G6PD levels in the HAMCs that can resist development of heme-free sGC in asthma. Hence the findings from this study places a new emphasis on the role of G6PD in lung sGC maturation in the airways that may potentially be a new target for treating asthma.

## MATERIAL AND METHODS

### Reagents

All chemicals were purchased from Sigma (St. Louis, MO) and Fischer chemicals (New Jersey). NO donor, NOC-18 or DETANONOate (2,2□-{Hydroxynitrosohydrazino) bis-ethanamine}), Phosphodiesterase inhibitor 3-isobutyl-1-methylxanthine (IBMX), sGC stimulator BAY 41-2272 (BAY-41, heme-dependent) and sGC activator BAY 60-2270 (BAY-60, heme-independent) were purchased from Sigma. Normal and patient-(severe asthma) derived HASMCs (de-identified and IRB exempt) were obtained from the Panettieri lab (Rutgers University) (Supplemental Table S1). Lung tissues from G6PD mice was obtained from Dr. Ling Wang (University of Iowa) (28). Cell lines, HEK293T and COS-7 were also purchased from American Type Culture Collection (ATCC; Manassas, VA, USA). Smooth muscle specific cell culture media was purchased from Lonza (Basel, Switzerland). Mammalian expression constructs for Myoglobin and G6PD were purchased from Origene (Rockville, MD, USA). Small interfering RNA (siRNA) specific to human G6PD (D-001830-01) was purchased from GE Healthcare Dharmacon (Lafayette, CO, USA). Real time PCR components including Trizol, cDNA kit, SYBR Green master mix etc. were obtained from Thermo Fisher Scientific and corresponding primers were obtained from Europhin. Protein G-sepharose beads were purchased from Sigma and molecular mass markers were purchased from Bio-Rad (Hercules, CA, USA).

### Antibodies

Antibodies were purchased from different sources. Supplemental Table S2 describes various types of antibodies used and its source.

### Cell culture, transfection and gene silencing by siRNA

All HASMCs were cultured in cell-specific media (Lonza), HEK or COS-7 cells were cultured and grown as previously described (4, 23). HASMCs were cultured to confluency (50-60%) and then treated with a low doses of NOC-18 (5 µM) for 16 h and then treated with BAY-41/60 for 10 min before cell harvest. Confluent non-NO treated cultures of both non-asthma/asthma HASMCs were also treated similarly with BAY-41/60 and supernatants generated. For HEK or COS-7 cultures, G6PD, Mb, sGCβ1 or sGCα1 were transfected for 36 h wherever applicable followed by −/+ NO or +SA/ +SA +hemin treatment. *Transfection/Co-transfection of G6PD SiRNAs:* For silencing of G6PD, 50 nM of human G6PD specific siRNA or scrambled siRNA (Dharmacon) was transiently transfected in select HASMCs for 48 h followed by 12 h of low NO treatment from NOC-18 before cell harvest.

### Western blots and Immunoprecipitations (IPs)

Cell supernatants or those that underwent transient transfections under −/+ NO or +SA / +SA +hemin treated conditions were analyzed for protein expression by western blots or for protein-protein interactions by immunoprecipitation assays (IPs). In other cases, mouse lung tissue segments obtained from naïve, OVA, CFA/HDM or G6PD mice lungs were processed to generate tissue supernatants for westerns. The asthma mice were generated by Dr. Kewal Asosingh’s lab as previously described (30). Band intensities on westerns were quantified using Image J quantification software (NIH).

### Quantitative real time PCR (qRT-PCR) analysis for mRNA expression in HASMCs

Cells from each NO treated or control groups (-/+NO) of HASMC (asthma and non-asthma) were harvested and processed for total RNA extraction using Trizol reagent. Total RNA was quantified using NanoDrop technology ND 1000 Spectrophotometer. Complimentary DNA (cDNA) synthesis was performed by reverse transcribing 200ng/µL of RNA using cDNA kit (Thermo Fisher Scientific, USA) according to the manufacturer’s protocol. Quantitative Real Time was carried out for measurement of *G6PD, TRX1, CYPB5R3*, and *sGC*β*1* mRNA expression on Biorad CFX96 Real-Time System using the cDNA prepared as described above. The following specific primers were used: *G6PD*-FP: ATG AGC CAG ATA GGC TGG AA, *G6PD*-RP: TAA CGC AGG CGA TGT TGT C, *TRX1*-FP: ATC AAG CCT TTC TTT CAT TCC CTC T, *TRX1*-RP: TTC ACC CAC CTT TTG TCC CTT C, *CYB5R3*-FP: GGA AGA TGT CTC AGT ACC TGG AG, *CYB5R3*-RP: TTG TCA GGT CGG ATG GCG AAC T, *sGC*β*1*-FP: ACA AGG AGA GGG CTG TAT CT, *sGC*β*1*-RP: AGA CGG AGG AAG GAC AGA ATA and β*-actin* FP: CGA CAA CGG CTC CGG CAT GTG C, β*-actin* RP: GTC ACC GGA GTC CAT CAC GAT GC.

### Immunostaining of human airway smooth muscle cells

Immunostaining was performed on HASMCs to determine the co-localization of sGCβ1 with G6PD under inactivated (-NO) condition in the two cell types (non-asthma or asthma) as previously described (22).

### Nitrite in the culture media

To assay for NO, nitrite was measured in the collected culture media using ozone-based chemiluminescence with the triiodide method and using the Sievers NO analyzer (GE Analytical Instruments, Boulder, CO, USA) (50).

### GSH/GSSG estimations and cGMP ELISA assays

Oxidized or reduced glutathione concentrations and the GSH/GSSG ratios in non-asthma or asthma HASMC supernatants were estimated using the GSH/GSSG assay kit (Bioassay Systems). For cGMP assays, the cell supernatants made from intact cells that were treated with NOC-18 were given BAY-41 before cell harvest, while others were treated with BAY-60 and their cGMP estimated by the ELISA assay kit (Cell Signaling Technology) (8).

## Acknowledgements

This work was supported by National Institute of Health Grants R56HL139564 and R01HL150049 (A.G.), Grants for R.A.P. (R01-HL166594), C.G. (R01NS137230) and D.J.S. (GM148664)

## Author Contributions

A. Ghosh, M. P. Sumi, C. Koziol-White, B. Tupta, L. Wang and C. Ghosh designed the experiments. A. Ghosh, M. P. Sumi, C. Koziol-White, B. Tupta, L. Wang and C. Ghosh performed all cell culture and biochemical studies. A. Ghosh, M. P. Sumi, C. Koziol-White, B. Tupta, L. Wang, C. Ghosh, W. F. Jester, R. A. Panettieri, and D. J. Stuehr analyzed all the data. A. Ghosh wrote the manuscript.

### Abbreviations

BAY-41: BAY 41-2272
BAY-60: BAY 60-2270
DEG: Differentially Expressed Gene
G6PD: Glucose 6-Phosphate Dehydrogenase
GSH: Glutathione
GSSG: Glutathione Disulfide
HASMCs: Human Airway Smooth Muscle cells
OVA: Ovalbumin
CFA/HDM: Complete Freunds Adjuvant/House Dust Mite
NOS: Nitric Oxide Synthase
PPP: Pentose Phosphate Pathway
PCLS: Precision Cut Lung Slices
PTMs: Post-translational Modifications
PAH: Pulmonary Arterial Hypertension
PASMCs: Pulmonary Arterial Smooth Muscle cells
ROS: Reactive Oxygen Species
SA: Succinyl Acetone

## Data Availability Statement Included in the article

The data that support the findings of this study are available in the methods, results and/or supplementary material of this article.

## Ethics Approval Statement

The present study followed international and/or institutional guidelines for humane animal treatment. The procedures involving animals were conducted on institutional guidelines and in compliance with Cleveland Clinic IACUC approved protocol.

