## Supplementary Information for "NO modulates human airway smooth muscle function by altering glucose-6-phosphate dehydrogenase effects on sGC function in asthma"

**Supplementary Information: Tables S1, S2, S3 and Figures S1-S7 with Figure legends**

**Supplementary Table S1. Donor information & use points in study**

**Supplementary Table S2. Antibodies used and their sources.**

**Supplementary Table S3. Summary of data obtained for each HASMC (Correlation Table).** Data are compiled using the results in the main or supplemental figures. The drug response listed for each HASMC indicates that a predominant sGC activation was achieved toward NO plus BAY-41 (NO + B41) or toward BAY-60. Protein expression levels, sGC heterodimer formation and sGC activation by NO + BAY-41 or BAY-60 are scored as Y(H/M): Yes (High/Moderate), Y(L): Yes (Low), ns/MR: non-significant/Muted response towards NO or ND: Not determined. *Criteria:* Low values are 1.5 to 4 fold less than High/Moderate values, while ns/MR are sometimes used for NO + B41 response or sGC heterodimer formation by low NO relative to the uninduced. These were determined for the 24 non-asthma or 25 asthma HASMC, and this information was then used to score that protein’s expression level, sGC heterodimerzation or activation in each non-asthma and severe asthma HASMC.

**Figure S1. Demographics representing race and gender distribution in all 40 non-asthma and 40 asthma HASMCs used in the study.**

**Figure S2.** **Impact of low NO from DETANONOate on sGC activation on asthma derived HASMCs.** Supernatants prepared from non-asthma or asthma HASMCs that were treated with -/+ low NO from DETANONOate as described in figure legend 2 were western blotted or determined for sGC activation (A) Expression levels as indicated and corresponding mean densitometries from n=16 non-asthma or n=17 asthma HASMCs, -/+ SD. (B) sGC activation by cGMP estimation on the ELISA as depicted in figure 2. *p < 0.05, by student’s unpaired *t*-test. Bars marked by * are significant relative to the non-NO treated HASMCs.

**Figure S3. Responders and non-responders to low NO.** HASMCs used from our earlier study (Redox Bio. 2021, Ref. 20) were cultured and treated with low NO from DETANONOate (16 h) plus BAY-41 before cell harvest. The supernatants were then assayed for G6PD expression by western blotting and their sGC activation estimated by cGMP measures. The Asthma HASMCs only partially responds to low NO, as 7 out of 10 asthma HASMCs showed a muted response to low NO. (A) Representative expression levels of G6PD as indicated. (B) Calculated densitometries of G6PD as depicted in panel A normalized to β-actin and calculated mean densitometries from non-asthma (n=16) and asthma (n=17) HASMCs -/+ SD. (C) cGMP estimated by ELISA. (D) Depicted linear correlation with mean cGMP values from sGC activation by NO+BAY-41 vs corresponding G6PD densitometries or expression levels. (Left) Tabulated sGC activation achieved toward NO plus BAY-41 (NO + B41) compared with G6PD densitometries for each individual HASMCs analyzed. G6PD expression levels and sGC activation by NO + BAY-41 are scored as Y(H/M): Yes (High/Moderate), Y(L): Yes (Low), MR: Muted response or ND: Not determined. *Criteria:* Low values are 1.5 to 4 fold less than High/Moderate values, while MR are sometimes used for NO + B41 response by low NO relative to the uninduced. These were determined for the 16 non-asthma or 17 asthma HASMC, and this information was then used to score the G6PD expression levels or sGC activation for indicated non-asthma and severe asthma HASMCs.

**Figure S4. Modulation of key protein expression by low NO in non-asthma or asthma HASMCs as depicted in figure 3.** To determine the molecular basis of the expression changes taking place at the protein level by low NO treatment, quantitative RT PCR was performed on a select group of some of the HASMCs. Following RNA extraction and cDNA synthesis on these HASMCs, specific primers were used to evaluate mRNA levels of sGCβ1, Cyb5r3 and Trx1 under conditions of -/+ NO treatment. (A) Densitometries of protein expression as shown in figure 3. (B) qPCR data on indicated genes from Set II depicting the impact of low NO on their corresponding mRNAs. CT values depicted are mean values, -/+ SD from n=3 independent experimental measures. *p < 0.05, by an independent student’s *t*-test, ns is not statistically significant. Bars depicted by asterisk (*) are significant relative to the other group.

**Figure S5. Correlation of G6PD densitometries or expression levels with corresponding GSH/GSSG ratios in non-asthma or asthma HASMCs as depicted in Fig. 4.**

**Figure S6. Immunostaining depicting co-localization of sGCβ1 and G6PD in non-activated (-NO) non-asthma or asthma HASMCs.** Non-activated (-NO) non-asthma or asthma HASMCs (15 & 15’ from Set II) were immunostained with sGCβ1 or G6PD antibodies. **(A**) Immunostained DAPI + sGCβ1 **(B**) Immunostained DAPI + G6PD **(C)** Immunostained images depicting sGCβ1 and G6PD co-localization. Images were captured at 10X with scale bar is 200 µM for all images.

**Figure S7. High NO and heme-free conditions influences G6PD expression.** COS-7 cells were either treated with SA for 48 h and transfected with sGCα1β1 or non-SA treated cells were transfected with sGCα1β1, G6PD or iNOS constructs (36 h) and as indicated certain combinations were treated with a low dose of NO donor from NOC-18 for 16 h before cell harvest. (A) Western blots depicting G6PD expression under normal, heme-free or overexpressed conditions -/+ NO treatment as mentioned in figure 5. (B) Corresponding mean densitometries of G6PD expression normalized to loading control β-actin as depicted panel A. Densitometries are mean (n=3 repeats) ± SD. *p < 0.05, by student’s *t*-test. Bar* 1 is significant to 2. ns is statistically non-significant.

**Supplementary Table S1. Donor information & use points in study.**

| **Redox Bio. 2021 study**  **(Ref. 20)** | **Pathology of**  **HASMCs** | **Age, Race, Gender** | **BMI** | **Tobacco use (Y/N)** | **Drugs, Medication taken, Cause of Death** | **Use points** |
| --- | --- | --- | --- | --- | --- | --- |
| 1 | Non-Asthma | 52/C/F | 28.8 | N | Nexium &topomax/  AVM malfunction/rupture | Fig. S3 |
| 2 | Non-Asthma | 17/C/M | 23.4 | Y | Marijuana/ GSW to head | Fig. S3 |
| 3 | Non-Asthma | 33/AA/F | 27.5 | Y | CVA, ICH | Fig. S3 |
| 4 | Non-Asthma | 50/C/F | 17.9 | N | Overdose of Insulin, Suicide | Fig. S3 |
| 5 | Non-Asthma | 22/AA/M | 25.1 | N | Anoxia, Blunt Injury | Fig. S3 |
| 6 | Non-Asthma | 55/C/M | 29.2 | N | ICH | Fig. S3 |
| 7 | Non-Asthma | 28/AA/F | 51.2 | N | CVA | Fig. S3 |
| 8 | Non-Asthma | 22/C/F | 23.1 | Y | SIGSW to Head | Fig. S3 |
| 9 | Non-Asthma | 22/C/F | 32.6 | Y | Anoxia, Seizure/MVA | Fig. S3 |
| 10 | Non-Asthma | 15/H/M | 14.5 | N | Marijuana/Adderall/  Head Trauma, GSW | Fig. S3 |
| 11 | Non-Asthma | 35/C/F | 26.7 | Y | CVA | Fig. S3 |
| 12 | Non-Asthma | 49/C/M | 38.7 | Y | CVA/Stroke | Fig. S3 |
| 13 | Non-Asthma | 29/C/M | 20.8 | Y | Marijuana /CVA, ICH | Fig. S3 |
| 14 | Non-Asthma | 53/H/M | 23.5 | N | CVA, ICH | Fig. S3 |
| 15 | Non-Asthma | 19/AA/M | 22.1 | Y | GSW | Fig. S3 |
| 16 | Non-Asthma | 13/H/F | 21.8 | N | MVA, Blunt injury | Fig. S3 |
| 1 | Asthma | 9/C/M | 11.2 | N | Albuterol /Asthma | Fig. S3 |
| 2 | Asthma | 22/AA/M | 36.3 | Y | Albuterol, Advair /Anoxia | Fig. S3 |
| 3 | Asthma | 22/AA/F | 23.0 | Y | Marijuana/Anoxia,  Asthma | Fig. S3 |
| 4 | Asthma | 46/C/F | 71.9 | N | Anoxia, Asthma | Fig. S3 |
| 5 | Asthma | 44/C/F | 34.5 | N | Respiratory Arrest | Fig. S3 |
| 6 | Asthma | 26/AA/F | 20.7 | N | Asthma Attack, Anoxia | Fig. S3 |
| 7 | Asthma | 54/C/M | 27.6 | N | Advair/Anoxia, | Fig. S3 |
| 8 | Asthma | 27/C/F | 30.4 | Y | Marijuana/Anoxia | Fig. S3 |
| 9 | Asthma | 35/C/F | 24.2 | N | Marijuana/Anoxia | Fig. S3 |
| 10 | Asthma | 19/C/F | 26.1 | Y | Albuterol/Asthma attack | Fig. S3 |
| 11 | Asthma | 38/C/M | 27.4 | N | Albuterol/Asthma attack | Fig. S3 |
| 12 | Asthma | 21/C/F | 43.9 | Y | Albuterol, Advair/Anoxia | Fig. S3 |
| 13 | Asthma | 14/H/M | 19.4 | N | Cardiac Arrest | Fig. S3 |
| 14 | Asthma | 14/C/F | 19.0 | Y | Advair/Cardiac Arrest | Fig. S3 |
| 15 | Asthma | 48/C/F | 26.7 | N | Heroin & Ecstay/ Advair, Albuterol/ Anoxia | Fig. S3 |
| 16 | Asthma | 12/AA/F | 21.4 | N | Singulair, Advair/Anoxia | Fig. S3 |
| 17 | Asthma | 44/H/F | 30.4 | N | Anoxia, CVA | Fig. S3 |
| **Current Study: Set I** | **Pathology of**  **HASMCs** | **Age, Race, Gender** | **BMI** | **Tobacco use (Y/N)** | **Drugs, Medication taken, Cause of Death** | **Use points** |
| 1 | Non-Asthma | 48/C/M | 32.79 | N | Drug Intoxication | Figs. 3, 4, 7, S2, S4, S5 |
| 2 | Non-Asthma | 22/B/M | 22.77 | Y: Cigarette/day x 1 yr | Head Trauma (GSW) | Figs. 3, 4, 7, S2, S4, S5 |
| 3 | Non-Asthma | 18/C/F | 24.11 | Y: ⅓ ppd  x 4 mth | Head Trauma (MVA) | Figs. 3, 4, 7, S2, S4, S5 |
| 4 | Non-Asthma | 43/H/M | 18.6 | Y: Unknown amount & duration | CVA/Stroke | Figs. 3, 4, S2, S4, S5 |
| 5 | Non-Asthma | 18/C/F | 30.85 | Y:Vape pen 10 puffs daily  x 1 yr | Head Trauma (MVA) | Figs. 3, 4, S2, S4, S5 |
| 6 | Non-Asthma | 41/C/F | 23.4 | Y:1 ppd  x 10 yrs | ICH/Stroke | Figs. 3, 4, S2, S4, S5 |
| 7 | Non-Asthma | 22/C/M | 20.36 | N | Head Trauma (GSW) | Figs. 2, 3, 4, 6, S2, S4, S5 |
| 8 | Non-Asthma | 24/B/F | 34.12 | N | Head Trauma (MVA) | Figs. 2, 3, 4, 6, S2, S4, S5 |
| 9 | Non-Asthma | 22/B/M | 24.36 | Y: Cigarette every 3 days  x 6 mth | Head Trauma (GSW) | Figs. 2, 3, 4, S2, S4, S5 |
| 10 | Non-Asthma | 20/C/M | 25.53 | Y:1pack/year x 12-15 mth | Head Trauma (MVA) | Figs. 2, 3, 4, S2, S4, S5 |
| 11 | Non-Asthma | 37/C/F | 22.41 | Y: ½ ppd x 21 yrs | CVA/Stroke | Figs. 2, 3, 4, S2, S4, S5 |
| 12 | Non-Asthma | 19/H/M | 25.19 | N | Head Trauma (MVA) | Figs. 2, 3, 4, S2, S4, S5 |
| 13 | Non-Asthma | 46/B/M | 28.56 | N | CVA/Stroke | Figs. 2, 3, 4, S2, S4, S5 |
| 14 | Non-Asthma | 17/H/M | 23.2 | N | Head Trauma (MVA) | Figs. 2, 3, 4, S2, S4, S5 |
| 1’ | Asthma | 14/C/M | 28.24 | N | Anoxia (Asthmatic Attack) | Figs. 3, 4, S2, S4, S5 |
| 2’ | Asthma | 22/C/F | 28.7 | N | Anoxia (Asthmatic Attack) | Figs. 3, 4, S2, S4, S5 |
| 3’ | Asthma | 26/C/F | 20.31 | N | Anoxia (Asthmatic Attack) | Figs. 3, 4, S2, S4, S5 |
| 4’ | Asthma | 11/B/M | 24.21 | N | Anoxia (Asthmatic Attack) | Figs. 3, 4, S2, S4, S5 |
| 5’ | Asthma | NA |  |  |  | Figs. 3, 4, S2, S4, S5 |
| 6’ | Asthma | NA |  |  |  | Figs. 3, 4, S2, S4, S5 |
| 7’ | Asthma | 44/C/F | 26.8 | N | Anoxia (Asthmatic Attack) | Figs. 2, 3, 4 6, S2, S4, S5 |
| 8’ | Asthma | 20/C/M | 20.5 | N | Anoxia (Asthmatic Attack) | Figs. 2, 3, 4, S2, S4, S5 |
| 9’ | Asthma | 27/B/M | 38.9 | Y: Cigarettes/  day x 7 mth | Anoxia (Asthmatic Attack) | Figs. 2, 3, 4, 7, S2, S4, S5 |
| 10’ | Asthma | 18/C/F | 24.6 | N | Anoxia (Asthmatic Attack) | Figs. 2, 3, 4, 6, S2, S4, S5 |
| 11’ | Asthma | 29/C/F | 35.01 | N | Anoxia (Asthmatic Attack) | Figs. 2, 3, 4, S2, S4, S5 |
| 12’ | Asthma | 25/B/F | 24.51 | N | Anoxia (Asthmatic Attack) | Figs. 2, 3, 4, 6, 7, S2, S4, S5 |
| 13’ | Asthma | 20/A/M | 26.83 | N | Anoxia (Asthmatic Attack) | Figs. 2, 3, 4, S2, S4, S5 |
| 14’ | Asthma | 37/C/F | 50.51 | Y:1-3 cigarettes/day x 20+yrs | Anoxia (Asthmatic Attack) | Figs. 2, 3, 4, 6, S2, S4, S5 |
| **Set (II)** | **Pathology of**  **HASMCs** | **Age, Race, Gender** | **BMI** | **Tobacco use (Y/N)** | **Drugs, Medication taken, Cause of Death** | **Use points** |
| 15 | Non-Asthma | 34/B/F | 54.06 | N | CVA/Stroke | Figs. 2, 3, 4, S4, S5, S6 |
| 16 | Non-Asthma | 19/H/M | 24.39 | N | GSW/Homicide | Figs. 2, 3, 4, 6, S4, S5 |
| 17 | Non-Asthma | 17/C/M | 22.1 | N | GSW/Homicide | Figs. 2, 3, 4, S4, S5 |
| 18 | Non-Asthma | 18/B/M | 21.76 | Y: 1-3 cigarettes/day x 2 yrs | Head Trauma (GSW) | Figs. 2, 3, 4, S4, S5 |
| 19 | Non-Asthma | 25/B/M | 32 | Y: 5-12 cigarettes/day x 4 yrs | Head Trauma (GSW) | Figs. 2, 3, 4, S4, S5 |
| 20 | Non-Asthma | 21/C/M | 21.97 | Y: less than 1 ppd x 4 yrs | Head Trauma (MVA) | Figs. 2, 3, 4, S4, S5 |
| 21 | Non-Asthma | 41/H/F | 30.27 | Y: 4 cigarettes/day x 3 yrs | ICH/Stroke | Figs. 2, 3, 4, S4, S5 |
| 22 | Non-Asthma | 20/C/M | 24.52 | N | Anoxia (Asphyxiation) | Figs. 2, 3, 4, S4, S5 |
| 23 | Non-Asthma | 12/C/F | 22.24 | N | Anoxia (Asphyxiation) | Figs. 2, 3, 4, 6, S4, S5 |
| 24 | Non-Asthma | 16/C/F | 20.42 | N | Head Trauma (MVA) | Figs. 2, 3, 4, S4, S5 |
| 15’ | Asthma | 30/B/M | 17.62 | N | Anoxia (Asthmatic Attack) | Figs. 2, 3, 4, S4, S5, S6 |
| 16’ | Asthma | 29/B/F | 89.12 | N | Anoxia (Asthmatic Attack) | Figs. 2, 3, 4, S4, S5 |
| 17’ | Asthma | 29/B/F | 29.89 | N | Anoxia (Asthmatic Attack) | Figs. 2, 3, 4, 6, S4, S5 |
| 18’ | Asthma | 11/B/M | 20.51 | N | Anoxia (Asthmatic Attack) | Figs. 2, 3, 4, S4, S5 |
| 19’ | Asthma | 41/B/F | 23.84 | N | Anoxia (Asthmatic Attack) | Figs. 2, 3, 4, S4, S5 |
| 20’ | Asthma | 13/B/M | 15.6 | N | Anoxia (Asthmatic Attack) | Figs. 2, 3, 4, 6, S4, S5 |
| 21’ | Asthma | 48/H/F | 36.27 | Y:Cigarette/  wk for < 1 yr | Anoxia (Asthmatic Attack) | Figs. 2, 3, 4, 6, S4, S5 |
| 22’ | Asthma | 18/C/M | 19.17 | N | Anoxia (Asthmatic Attack) | Figs. 2, 3, 4, S4, S5 |
| 23’ | Asthma | 11/C/M | 30.61 | N | Anoxia (Asthmatic Attack) | Figs. 2, 3, 4, S4, S5 |
| 24’ | Asthma | 23/H/M | 20.76 | N | Anoxia (cardiovascular) | Figs. 2, 3, 4, S4, S5 |
| 25’ | Asthma | 21/C/F | 43.9 | Y: 4 cigarettes/day x 5 yrs | Anoxia (Asthmatic Attack) | Figs. 2, 3, 4, S4, S5 |

Abbreviations: C; Caucasian, H; Hispanic, F; Female, M; Male, AA; African American, CVA; Cerebrovascular accident, AVM; Arteriovenous malformation, ICH; Intra-cerebral hemorrhage, SIGSW; Self inflicted gunshot wound, GSW; Gunshot wound, MVA; Motor vehicle accident, Unk; Unknown. NA; Not Available. The HASMC are numbered as depicted in the leftmost part of the Table and these numbers are used in the manuscript.

**Supplementary Table S2. Antibodies used and their sources.**

| **Serial No.** | **Antibody** | **Species-Clonality** | **Source (Catalog No.)** |
| --- | --- | --- | --- |
| 1. | sGCβ1 | Rabbit polyclonal | SIGMA, ER-19 (G4405) |
| 2. | sGCβ1 | Rabbit polyclonal | SIGMA, S-15 (G4530) |
| 3. | sGCβ1 | Rabbit polyclonal | Cayman Chem. (160897) |
| 4. | sGCα1 | Rabbit polyclonal | Abcam (ab50358) |
| 5. | sGCα1 | Rabbit polyclonal | Abcam (ab101368) |
| 6. | sGCα1 | Rabbit polyclonal | Cayman Chem. (160895) |
| 7. | G6PD | Mouse monoclonal | Santa Cruz (sc-373886) |
| 8. | G6PD | Mouse monoclonal | Santa Cruz (sc-373887) |
| 9. | Hsp90 | Mouse monoclonal | Origene (TA500494) |
| 10. | Hsp90 | Rabbit polyclonal | Cell Signaling Tech. (4875S) |
| 11. | β-Actin | Mouse monoclonal | SIGMA (A5441) |
| 12. | CYB5R3 | Mouse polyclonal | Abnova (H00001727-A01) |
| 13. | Myc-tag | Mouse monoclonal | Cleveland Clinic |
| 14. | Myc-tag | Rabbit monoclonal | Cell Signaling Tech. (2278S) |
| 15. | V5-tag | Mouse monoclonal | Thermo Fischer (46-0705) |
| 16. | Catalase | Mouse monoclonal | Santa Cruz (sc-271803) |
| 17. | Trx1 | Rabbit polyclonal | Origene (TA322865) |
| 18. | Trx1 | Rabbit monoclonal | Cell Signaling Tech. (2429S) |

**Supplementary Table S3. Summary of data obtained for each HASMC.**

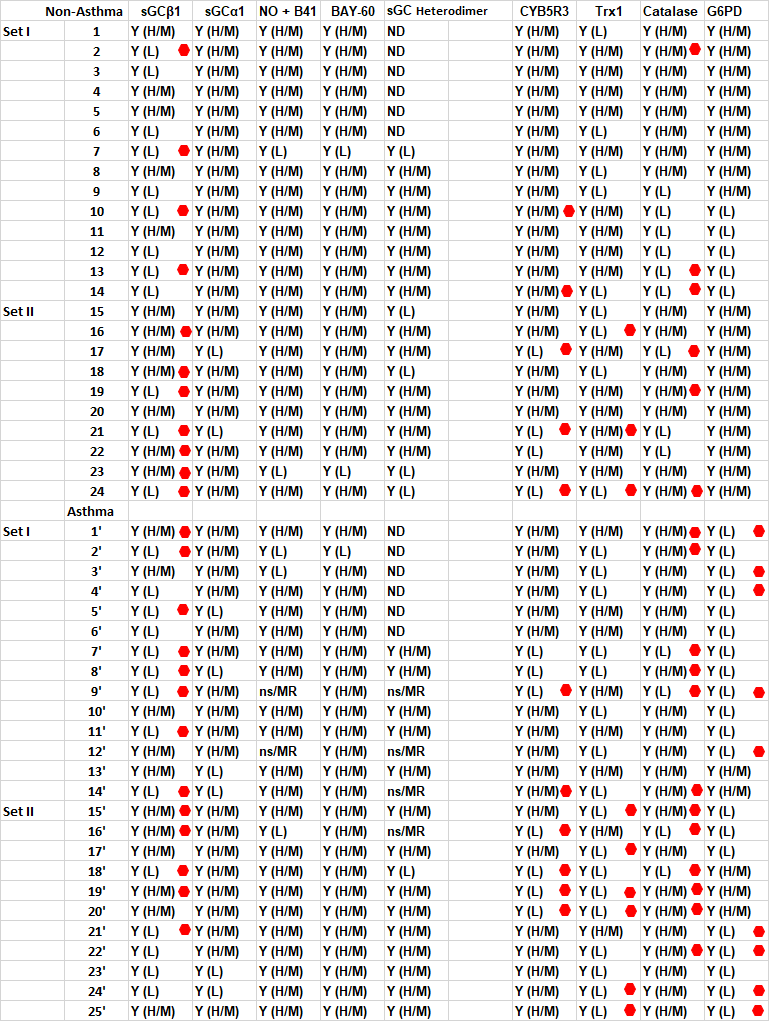

Data are compiled using the results in the main or supplemental figures. The drug response listed for each HASMC indicates that a predominant sGC activation was achieved toward NO plus BAY-41 (NO + B41) or toward BAY-60. Increase in protein expression by low NO is denoted by . Protein expression levels, sGC heterodimer formation and sGC activation by NO + BAY-41 or BAY-60 are scored as Y(H/M): Yes (High/Moderate), Y(L): Yes (Low), ns/MR: non-significant/Muted response towards NO or ND: Not determined. *Criteria:* Low values are 1.5 to 4 fold less than High/Moderate values, while ns/MR are sometimes used for NO + B41 response or sGC heterodimer formation by low NO relative to the uninduced. These were determined for the 24 non-asthma or 25 asthma HASMC, and this information was then used to score that protein’s expression level, sGC heterodimerzation or activation in each non-asthma and severe asthma HASMC.

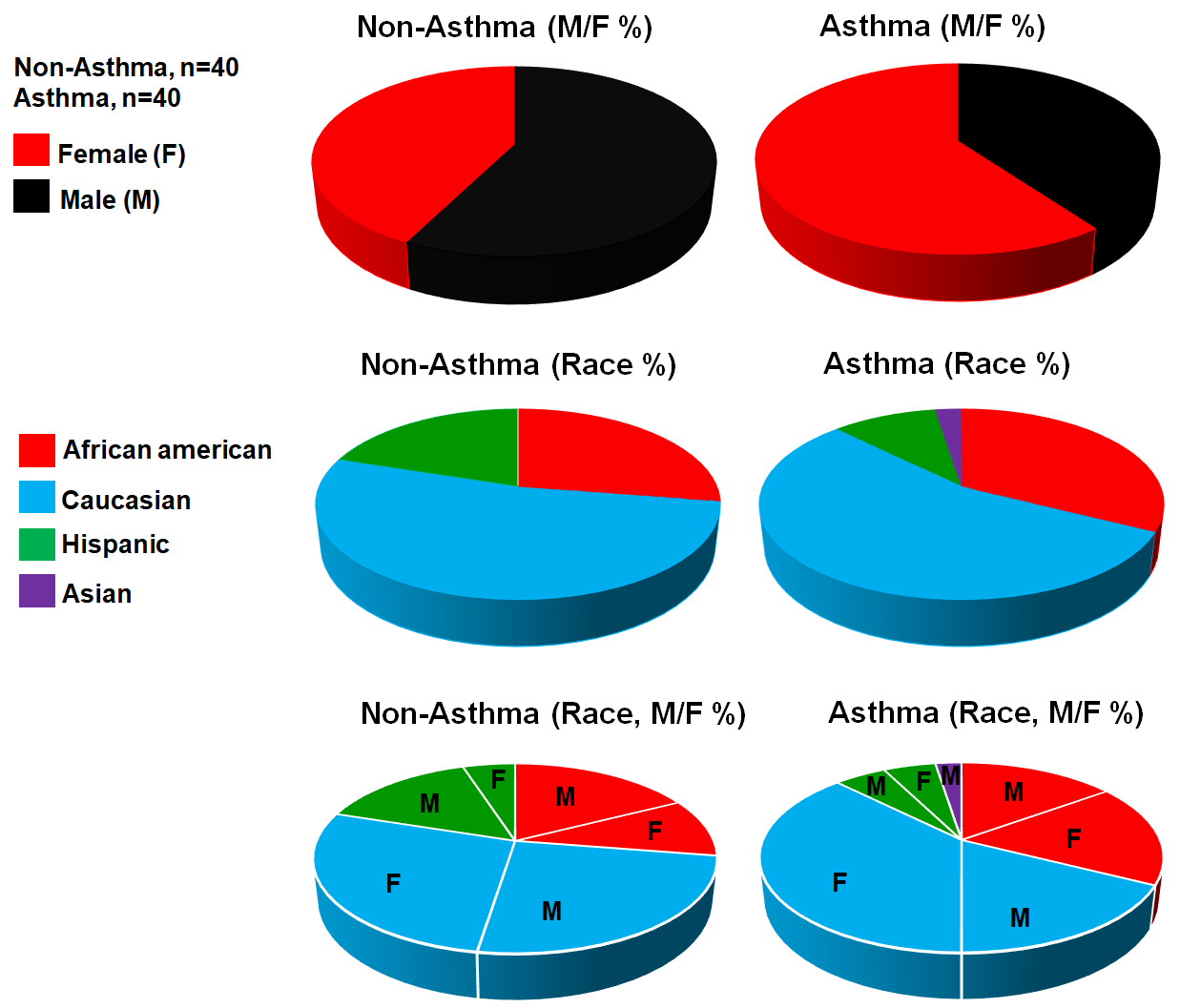

**Figure S1. Demographics representing race and gender distribution in all 40 non-asthma and 40 asthma HASMCs used in the study.**

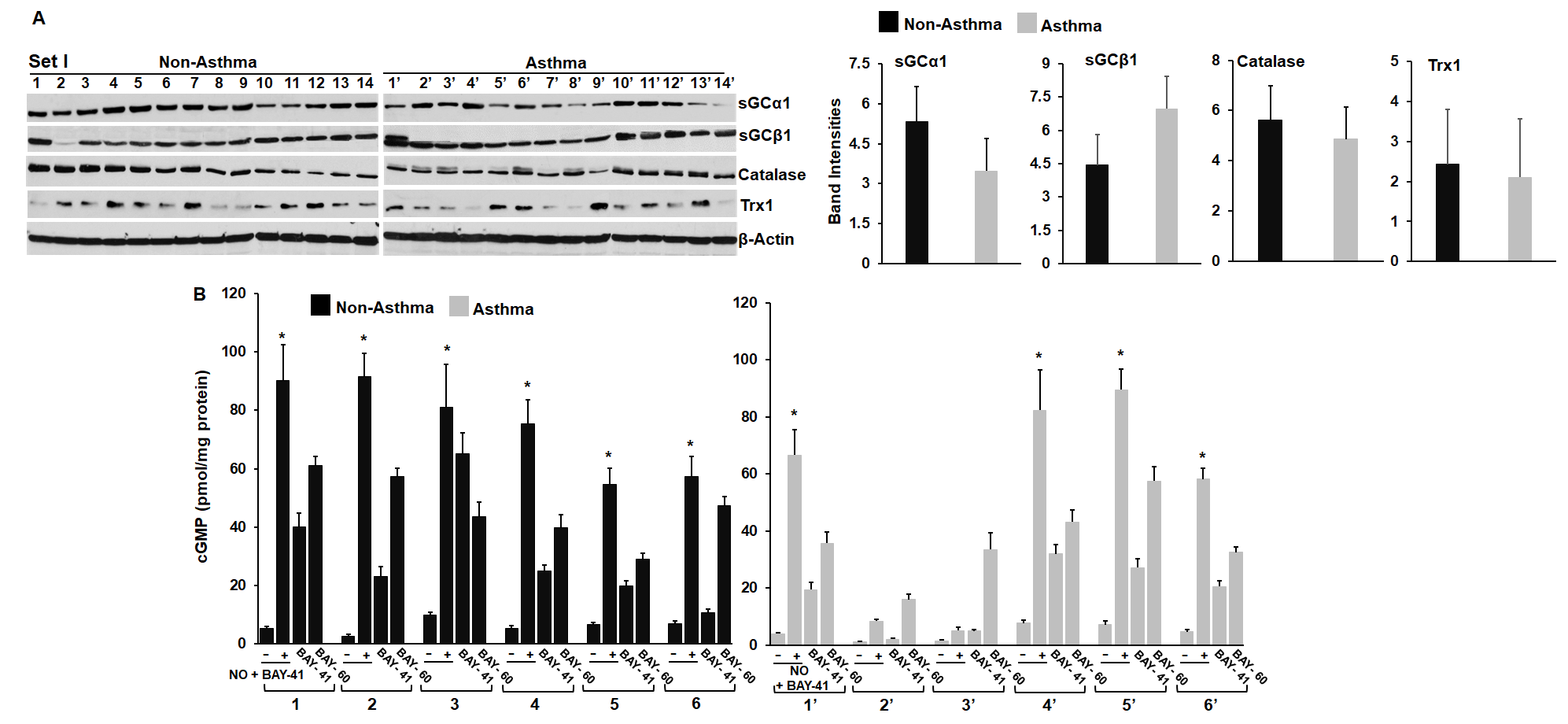

**Figure S2.**

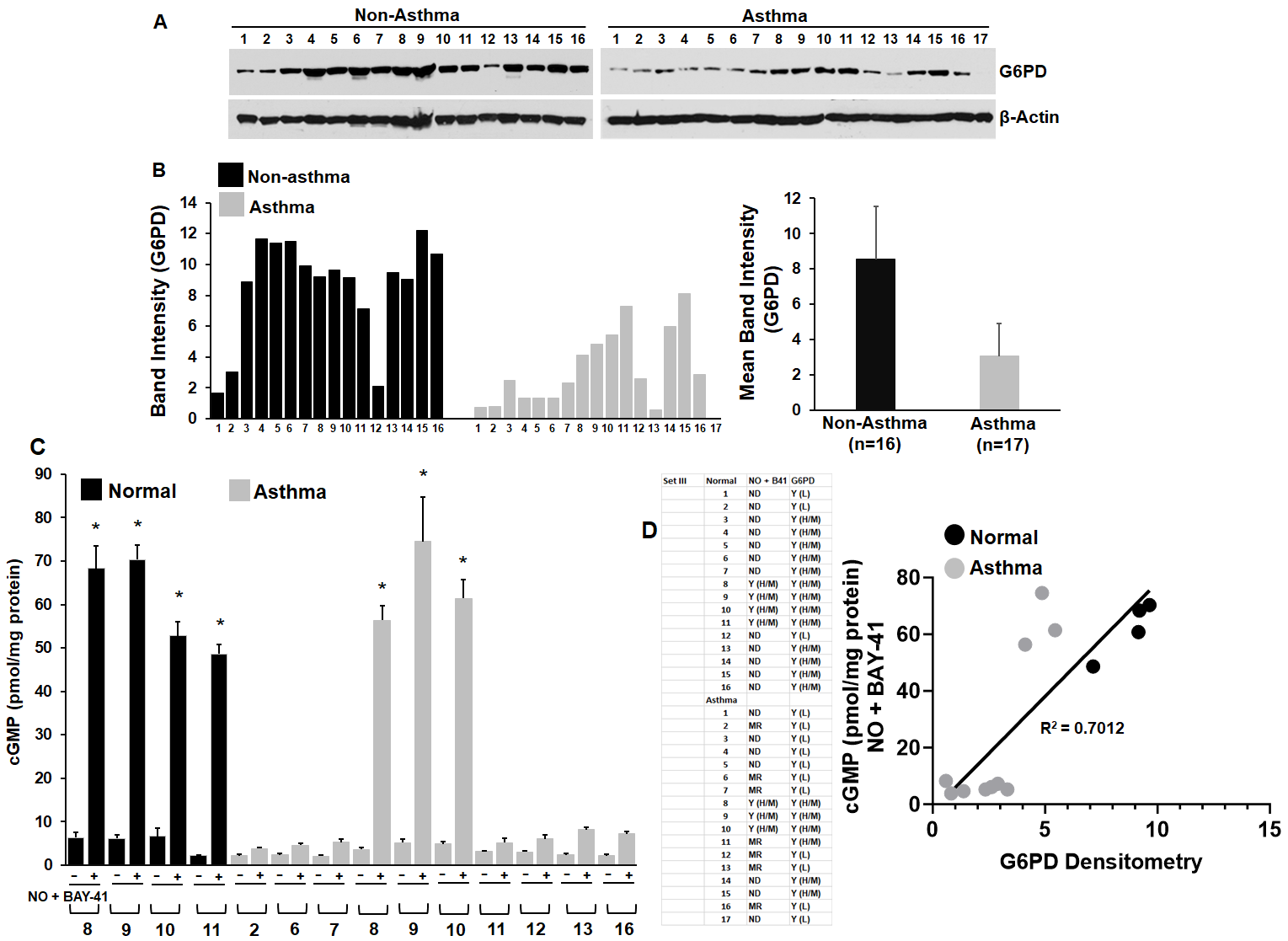

**Figure S3.**

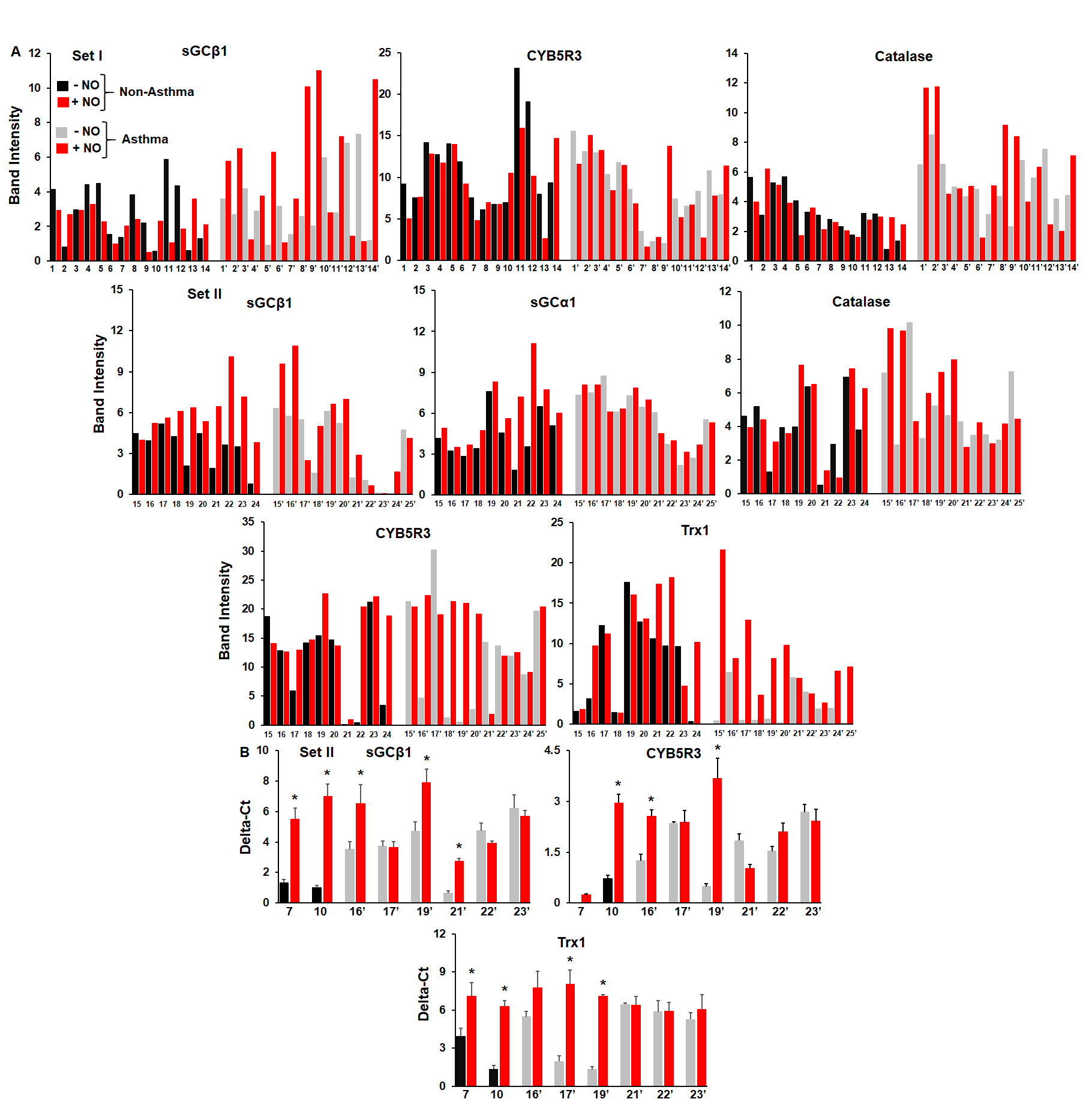

**Figure S4.**

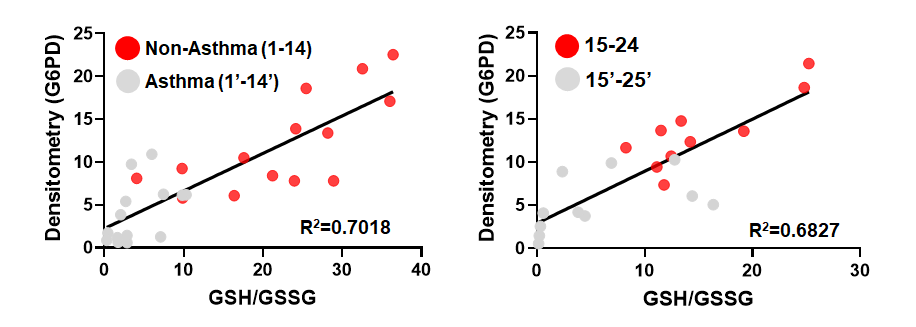

**Figure S5. Correlation of G6PD densitometries or expression levels with corresponding GSH/GSSG ratios in non-asthma or asthma HASMCs as depicted in Fig. 4.**

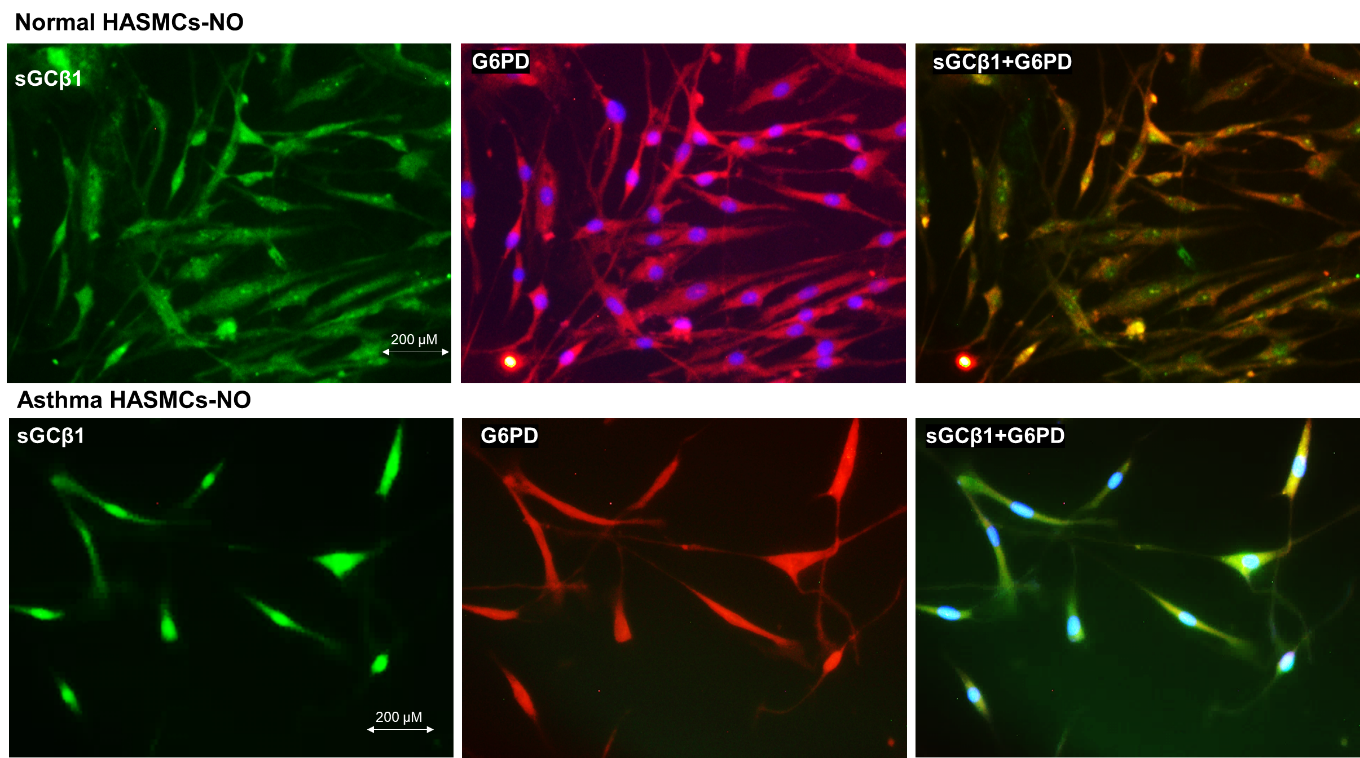

**Figure S6.**

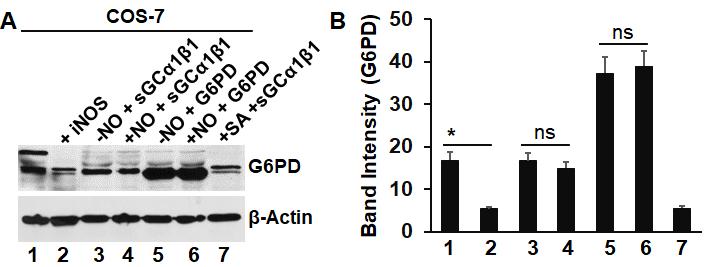

**Figure S7.**
